# Stomatal and xylem plasticity, not growth rate, determines white spruce resilience to warmer and drier climates

**DOI:** 10.64898/2026.09.02.748918

**Authors:** Bridget K. Murphy, Noelle Perkins, Fatima Nagi, Siyu Wang, Tyler Muchos, Julia A. Boyle, Nathalie Isabel, Ingo Ensminger

**Author notes:** **Corresponding author**: Ingo Ensminger.

## Abstract

In a warmer and drier climate, forest productivity will depend on trees’ ability to maintain carbon uptake and hydraulic function. Whether fast-growing genotypes of boreal conifers are more vulnerable to combined climatic stress remains uncertain. Using a full-factorial field experiment, we investigated how progressive soil drying combined with extended warming affects growth, xylem development, and photosynthesis in two *Picea glauca* families with contrasting growth strategies. Rainout structures first reduced soil moisture from 25% to 18%, followed by a +5 °C warming treatment applied using infrared heaters. During the warmest and driest period in August, air temperature reached 34.5°C in the warmed plots, while soil moisture declined to a low of 15% in the combined rainout and warming treatment. Contrary to expectations, both fast- and slow-growing white spruce families exhibited similar resilience to concurrent warming and soil drying. This finding challenges the prevailing theory that faster growth increases vulnerability to climatic stress. Despite an approximately 50% reduction in rainfall, pre-dawn water potential remained above –0.5 MPa across treatments, reflecting that seedlings were able to avoid hydraulic stress. Although the fast-growing family maintained greater height and diameter growth compared to the slow-growing family, both exhibited similar physiological and anatomical responses to warming. Warming decreased stomatal conductance, which increased intrinsic water-use efficiency. Latewood xylem traits related to hydraulic efficiency were also reduced under warming. Together, these coordinated stomatal and xylem adjustments decreased water loss and protected hydraulic function, enabling both families to maintain high photosynthesis and growth under simulated climate conditions. Overall, white spruce exhibits strong phenotypic plasticity, supporting intraspecific resilience to moderate warming and soil drying representative of projected 21st-century summer conditions for central and eastern Canada.

## Introduction

Heatwaves are increasing in frequency, intensity and duration globally due to climate change (Rahmstorf & Coumou, 2011). In 2021, North America experienced one of the most extreme heatwaves on record, with temperatures exceeding climatological norms by up to 16°C (Thompson *et al*., 2022; Baum *et al*., 2026). Comparable record-breaking events have also been documented in Europe (García-Herrera *et al*., 2010), Asia (Pattanaik *et al*., 2017), and Australia (Coates *et al*., 2014). Heatwaves reduce primary productivity (Ciais *et al*., 2005; Xu *et al*., 2020; Zhu *et al*., 2021), increase wildfire risk (Hegedűs *et al*., 2024), and threaten human health (Coates *et al*., 2014). The IPCC defines a heatwave as at least five consecutive days with maximum temperatures at least 5°C higher than the climatological norm. Yet most experimental studies simulate more extreme conditions, often exceeding control temperatures by 10°C or more (Duarte *et al*., 2016; Guha *et al*., 2018; Birami *et al*., 2018; Dhami *et al*., 2020; Gagne *et al*., 2020; O’Connell & Wiley, 2024), with some including a “moderate” heatwave treatment 6°C above control (Ameye *et al*., 2012; Bauweraerts *et al*., 2014). Heatwaves often coincide with low precipitation, prompting recent studies to simulate more realistic experimental designs that combine warming and water limitation (De Boeck *et al*., 2010; Bauweraerts *et al*., 2014; Drake *et al*., 2018; Birami *et al*., 2020; Gagne *et al*., 2020; Marchin *et al*., 2022).

Under combined warming and drought, plants face a physiological trade-off between evaporative cooling and water conservation to avoid hydraulic failure. When water is limited, plants typically reduce stomatal conductance to limit transpiration, restricting CO_2_ uptake for carboxylation by Rubisco and thereby suppressing photosynthesis (Joshi *et al*., 2022). Reduced transpiration also impairs evaporative cooling, increasing leaf temperatures and intensifying heat stress (Farooq *et al*., 2009; Feller, 2016). During heatwaves, drought-treated trees can exhibit leaf temperatures 2-3°C higher than well-watered trees (Birami *et al*., 2018). Even without drought, heat stress alone decreases rates of photosynthesis, damages photosystem II (PSII), decreases electron transport, hampers thylakoid membrane integrity, and inactivates Rubisco (Wahid *et al*., 2007; Teskey *et al*., 2015). High temperatures also elevate respiration rates and hence increase loss of previously fixed carbon (Atkin & Tjoelker, 2003). When photosynthesis declines while respiration rises, the carbon balance can shift with more carbon being respired than fixed. Drought often amplifies this imbalance. For example, in white cedar (*Thuja occidentalis)*, warming caused greater carbon losses under drought because respiration increased linearly with increasing temperature whereas CO₂ assimilation declined earlier under mild drought (Zhao *et al*., 2013).

These physiological constraints require trees, including needle-leaved conifers, to deploy photoprotective strategies that safeguard photosynthetic machinery and dissipate excess light energy. When photosynthesis is inhibited, conifers upregulate carotenoids, such as lutein and *β*-carotene, and rely on non-photochemical quenching (NPQ; Ensminger *et al*., 2004; Demmig-Adams *et al*., 2012; Fréchette *et al*., 2016; Murphy *et al*., 2025). NPQ, measurable via chlorophyll fluorescence, is driven by rapid conversions of xanthophyll cycle pigments in which violaxanthin is de-epoxidated into antheraxanthin and zeaxanthin. Carotenoid-based vegetation indices, such as the photochemical reflectance index (PRI) and the chlorophyll carotenoid index (CCI), can detect stress-related changes in foliar pigment composition correlated with photosynthetic downregulation (D’Odorico *et al*., 2021; Murphy *et al*., 2025). These indices can be measured non-destructively using leaf spectral reflectance, enabling monitoring of climatic-stress resilience and scaling through remote-sensing. The water index (WI), which captures changes in the near-infrared spectrum, has also been used to effectively monitor leaf water content and soil-moisture effects in some tree species (Claudio *et al*., 2006; Junttila *et al*., 2015).

Beyond leaf-level photoprotection, structural traits, such as xylem anatomy, are critical for sustaining water transport under concurrent heat and drought. Drought alone can induce xylem embolism and reduce hydraulic conductivity (Lewis, 1988; Tyree & Sperry, 1988; McDowell *et al*., 2008), and the addition of heat further exacerbates hydraulic strain by driving more negative water potentials and reducing leaf hydraulic conductance by as much as 90% (Rehschuh & Ruehr, 2022). Conifers respond to these challenges through a combination of avoidance, resistance, and recovery (Baldi & La Porta, 2022). Among these strategies, a plant’s capacity to endure water stress through constitutive or induced traits, such as xylem architecture, is critical to maintaining function under compound stress. Conifer species resistant to xylem cavitation often possess thicker tracheid cell walls with only modest reductions in lumen diameter, helping preserve conductivity while reducing risk of hydraulic failure (Bouche *et al*., 2014). However, the effectiveness of such structural adjustments can vary across individuals and environments. Phenotypic plasticity in hydraulic and photosynthetic traits presents an opportunity for conifers to cope with increasing climate variability (Aubin *et al*., 2016; Ravn *et al*., 2022), although the extent to which this plasticity enhances resilience under future warming and drying remain uncertain.

Most studies of conifer responses to combined heat and drought focus on species-specific patterns, overlooking substantial intraspecific variation among genotypes and families. Dendroecological evidence for intraspecific variation in conifer drought responses comes mostly from studies of mature, established trees (Zas *et al*., 2020; Schueler *et al*., 2021; Depardieu *et al*., 2024), leaving seedlings, despite their critical vulnerability and crucial role in reforestation, underrepresented. As a result, xylem development and plasticity during the seedling stage remains poorly understood. A recent global dendroecological study on mature conifer species suggest a trade-off exists between fast- and slow-growth strategies (Wang & Wang, 2024), consistent with the ‘fast–slow’ plant economics spectrum theory that characterizes plants along a continuum from resource-acquisitive to resource-conservative strategies (Reich, 2014). Along this spectrum, fast-growing species tend to exhibit resource-acquisitive traits, such as higher maximum rates of photosynthesis and specific leaf area, while slow-growing species typically possess more conservative traits geared toward stress tolerance, such as higher wood density (Reich, 2014). In line with this theory, drought-tolerant conifers typically exhibit higher wood density, greater hydraulic safety margins, more negative P_50_ values (the water potential at which 50% of hydraulic conductivity is lost), shorter maximum tree height and lower specific leaf area (Wang and Wang 2024). This raises an important question: are fast-growing genotypes more vulnerable to heat and drought stress than those with slower growth?

Evidence from common-garden and growth-chamber experiments supports this hypothesis. In a common garden, white spruce populations originating from drier climates maintained greater radial growth during severe droughts in 2001-2002 compared to populations from more humid climates (Depardieu *et al*., 2020). Although such studies reveal genetic variation in drought tolerance, considerably less is known about how warming and drought interact to affect populations with different growth strategies. In growth chambers, severe drought (water potential, WP < −2.0 MPa) and extreme short-term warming to 42-50°C individually had stronger negative effects on high-growth white spruce families than on intermediate- or slow-growth families (Bigras, 2000, 2005). Under more moderate drought (WP > −2.0 MPa), however, families responded similarly, suggesting sensitivity depends on stress severity. The heat-shock temperatures used greatly exceeded those typically experienced in natural environments. In Canada, heatwaves in 2024-2025 reached maximum temperatures of 30.9°C (Environment and Climate Change Canada, 2025), far below the 42–50°C applied experimentally. This discrepancy highlights a key gap: while fast-growing families appear disadvantaged under severe stress, their responses to moderate but ecologically realistic warming, especially when combined with drought, remain unclear.

Addressing this gap requires moving beyond controlled environments and evaluating conifer responses under conditions that more closely mirror natural heatwaves. Accordingly, we designed a factorial field experiment to examine how photosynthesis, respiration, growth and water relations of a fast- and a slow-growing white spruce family respond to progressive soil drying concurrent with warming. We hypothesized that: (i) fast-growing families show similar resilience comparable to slow-growing families under moderate single stressors, but reduced resilience when warming and soil-moisture deficit co-occur; and (ii) photosynthesis and growth decline more under combined warming and soil-moisture deficit than under either stressor alone. In this study, resilience denotes the capacity to maintain functional, hydraulic, and structural performance during climatic stress and was assessed through a range of physiological, growth, and xylem anatomical measurements.

## Materials and Methods

### Field Site and Plant Materials

The experiment was conducted at the University of Toronto’s Koffler Scientific Reserve, near King City, ON (44°01’44.3”N 79°32’19,2”W). Plots were excavated and filled with equal parts peat, sand, and local soil (New Roots Garden Center). One-year-old bare-rooted white spruce (*Picea glauca* [Moench] Voss) seedlings were provided by Natural Resources Canada (Laurentian Forestry Centre), planted in 2017 and were five years old at experiment onset. Two full-sib families were selected from the ten planted, as they displayed significant differences in initial stem height and diameter, indicative of different growth strategies. One family originated from a controlled cross of two Peterborough, Ontario genotypes [44°18’17” N, 78°19’12” W], and the other from a cross of two Sundridge, Ontario genotypes [45°46’08”N, 79°23’47”W] (Fig. **1**). Climate conditions at these parental sites are shown in Fig. **S1**. Six seedlings per family were planted per plot at ∼25 cm spacing with an outer ring of buffer seedlings to mitigate edge effects. Plots were ∼1 m apart. Additional planting and plot maintenance details are provided in Murphy *et al*. (2025).

**Figure 1.**
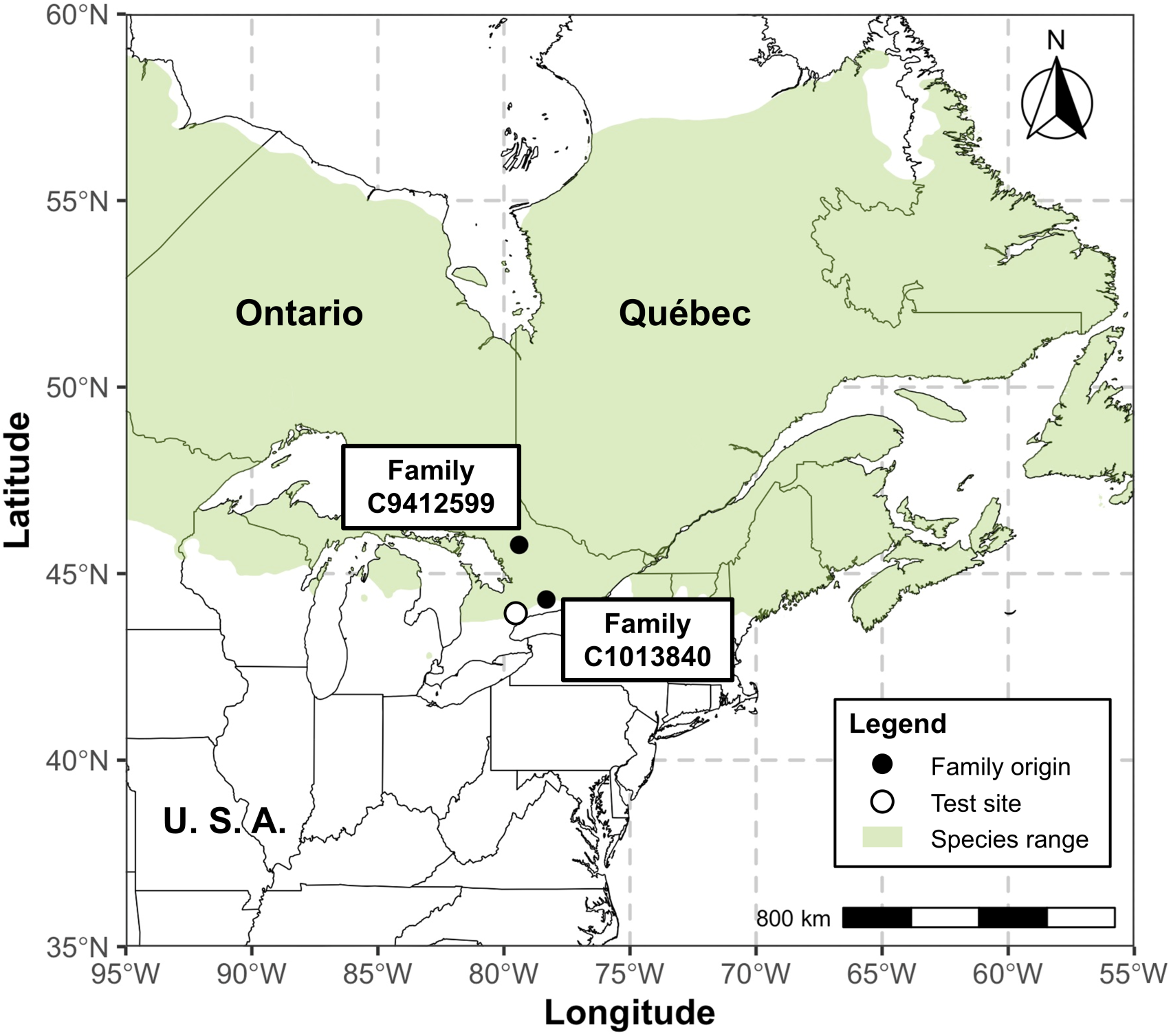
A map displaying the parental origins of the two families planted in a common garden experiment at Koffler Scientific Reserve in King City, Ontario, Canada. Family C9412599 is a cross between two parents from Sundridge, Ontario that has been characterized as slow-growing. Family 1013840 is a cross between two parents from Peterborough, Ontario, that has been characterized as fast-growing. Growth strategy differences were assessed based on height and diameter measurements taken in May, prior to the start of the experiment. Political boundaries for both Canada and the U.S. were obtained from Natural Earth (Natural Earth, 2025). White spruce (*Picea glauca*) species range polygons were obtained from digitized shapefiles available on DataBasin, based on Little, E.L., Jr., 1971, Atlas of United States Trees, U.S. Department of Agriculture, Forest Service.

To induce a soil-moisture deficit concurrently with warming, treatments were introduced sequentially. First a rainout (RO) treatment was used to exclude rainfall and reduce soil moisture, followed by warming (W), applied either alone or combined with RO (RO+W; Fig. **S2**). A fourth treatment served as the ambient control. RO structures were installed on June 24 and warming began on Aug 16, six weeks after RO installation (Fig. **2**). Each treatment had three plot replicates. Soil water content was reduced from late June to late September using RO structures, and leaf temperature was increased by 5°C during night and day in warming plots using infrared heaters from mid-August to late September. Technical details on the Temperature Free-Air-Controlled Enhancement (T-FACE) system are provided in Frechette *et al*. (2020). For the RO treatment, six plots had rainout structures with open profiles (longitudinally halved PVC pipes) installed between the seedlings at an incline to intercept ∼ 50% of the precipitation and divert it via gutters and tubing away from all experimental plots. Control non-RO plots had inverted “dummy” profiles installed that allowed rainfall to run down and drip into the soil. To prevent unintended drought, control plots were watered to maintain a targeted soil moisture of 25%.

**Figure 2.**
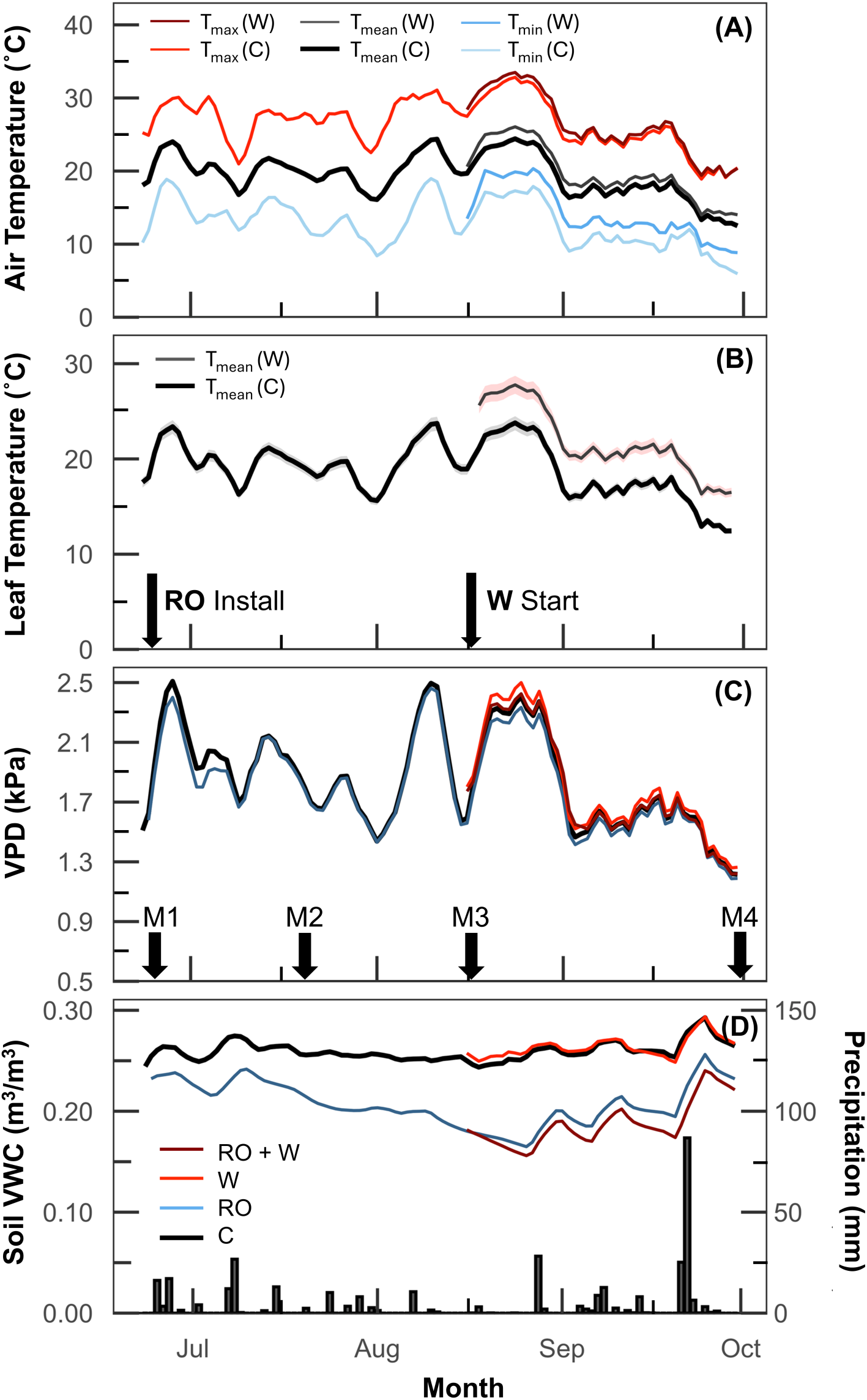
Seasonal variation in environmental and experiment conditions from June 24 to October 1, 2021, at the Koffler Scientific Reserve in Ontario, Canada. **(a)** 5-day running averages of daily minimum (T_min_), mean (T_mean_), and maximum (T_max_) air temperature in warming (W) and control (C) plots; (**b)** 5-day running averages of estimated daily mean leaf temperatures (T_mean_) in warming (W) and control (C) plots, with 95% confidence intervals; (**c**) 5-day running average of daily vapour pressure deficit (VPD); **(d)** 5-day running average of daily soil volumetric water content (VWC) across treatments and daily total precipitation (mm). Control, C; Rainout, RO; Warming, W; Rainout combined with warming, RO+W. Black arrows represent the installation of rainout structures and the onset of warming, and white arrows indicate measurement (M) campaigns. Sampling was limited to M3 and M4.

Each plot was equipped with a canopy-level air temperature and humidity sensor (Models VP3 and VP4, METER Group), and a soil temperature and moisture sensor at a depth of ∼20cm (Models 5TM and TE11, METER Group), logging data every 30-minutes using a CR1000 datalogger (Fig. **2**; Method S1; Campbell Scientific Inc.). Air temperature was used to estimate leaf temperature due to non-operational plot-level leaf temperature sensors, with estimates derived from regression models trained on air and leaf temperature data collected in 2020 under the same experimental design (Murphy *et al*. 2025; Method S2). A weather station recorded site-wide conditions every 15 minutes with an air temperature and humidity sensor (Model Rotronic HC2A-S3, ITM Instruments), a PAR sensor (LI-COR LI-190R Quantum Sensor, Li-Cor Biosciences), a solar flux sensor (Model SP Lite2 Pyranometer, OTT HydroMet), and a tipping bucket rain gauge (Model 52202, R.M. Young Company). Occasional gaps in precipitation data were supplemented using records from the King City North weather station, 8.5 km from the field site (Environment Canada).

The experiment ran from June 23 to October 1, 2021, spanning 101 days. Four measurement campaigns were conducted on days 1 (prior to any treatment), 31 (∼4 weeks after onset of RO), 54 (∼7 weeks after onset of RO) and 98 (∼6 weeks after onset of warming with existing RO). Each campaign included measurements of photosynthetic gas exchange, chlorophyll fluorescence, spectral reflectance, growth, and water potential. Destructive needle sampling for pigment analysis was performed only twice on days 31 and 98 to minimize damage of seedlings and conserve tissue. Most campaigns spanned two consecutive days: physiological measurements between ∼8:00-17:00 hr on day one, followed by sample collection after pre-dawn water potential at 4:00 hr on day two. The final campaign required three days to accommodate additional gas exchange measurements. Chlorophyll fluorescence, spectral reflectance, and tissue sampling for xylem analyses were performed using two seedlings per family, averaged to calculate the plot average for each family. Each plot average was used as a biological replicate, yielding three replicate plots per treatment. Photosynthetic gas exchange, water potential measurements, and tissue sampling for pigment analyses was performed using one seedling per family.

### Chlorophyll Fluorescence Measurements

Chlorophyll fluorescence was measured using a Dual-PAM-100 fluorometer (Walz). A bundle of attached, current-year sunlit needles was arranged as a flat layer and placed into a dark leaf clip (DLC-8, Walz), then dark acclimated for 30 minutes prior to measurement. Minimal fluorescence (F_o_) was recorded, followed by a short saturation pulse to determine maximal fluorescence (F_m_) of the dark acclimated leaves. Maximum quantum yield of PSII (F_v_/F_m_) was calculated according to Genty *et al*. (1989):

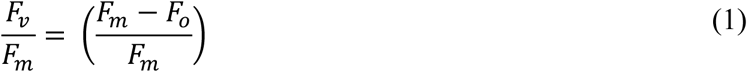

Needles were then exposed to 1,500 μmol photon m^-2^ s^-1^ for five minutes to induce steady-state fluorescence (F). A weak far-red light pulse was applied to determine minimal fluorescence of light adapted needles (F_o_′), followed by a saturating pulse to measure maximum fluorescence (F_m_′). Excitation energy partitioning was estimated according to Hendrickson *et al*. (2004), where effective quantum yield (ϕ_PSII_) represents the light energy that is used for photochemistry, dynamic non-photochemical quenching (ϕ_NPQ_) represents the light energy that is being dissipated as heat by the xanthophyll cycle pigment pool, and sustained non-photochemical quenching (ϕ_f,D_) represents constitutive energy dissipation. Excitation pressure on PSII (1-qP) is a measure of the redox state of the Q_A_ pool, represented by Equation 5, and was calculated according to Huner *et al*. (1998)

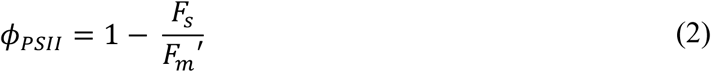

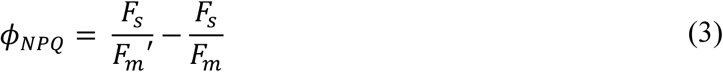

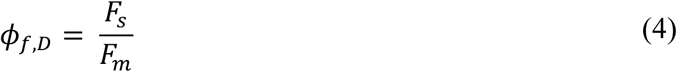

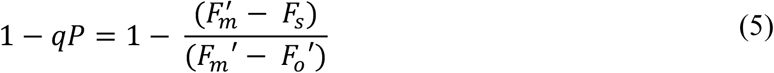

### Leaf Spectral Reflectance Measurements

Leaf spectral reflectance was measured with a portable spectrometer (UniSpec-SC, PP Systems) using a bifurcated fibre-optic (UNI410, PP Systems) and a needle leaf clip (UNI501, PP Systems). Measurements were taken at a fixed 60° angle relative to the needle axis. A dark-current correction scan preceded white reference scans. Integration time was set to 10 ms, with automatic averaging of 10 scans. The Photochemical Reflectance Index (PRI), a carotenoid-based vegetation index that tracks photosynthetic efficiency through shifts in xanthophyll cycle pigment pools relative to chlorophyll, was calculated according to Peñuelas et al. (1995):

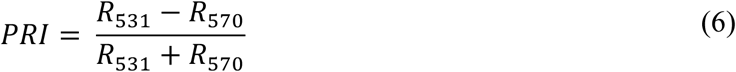

where R_531_ and R_570_ indicates reflectance at 531 and 570 nm, respectively. High PRI values indicate photosynthetically favorable performance, where light is efficiently used for photochemistry while excess energy is dissipated through the xanthophyll cycle (Wong & Gamon, 2015). The Chlorophyll Carotenoid Index (CCI), which tracks seasonal changes in leaf carotenoid pigment pools relative to chlorophyll, was calculated according to Gamon et al. (2016):

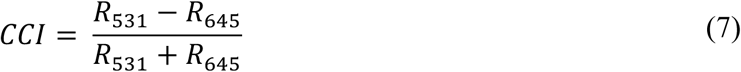

where R_531_ and R_645_ indicates reflectance at 531 and 645 nm, respectively. High CCI values indicate higher chlorophyll content relative to carotenoids, reflecting healthy foliage and robust photosynthetic activity (Gamon *et al*., 2016). The normalized vegetation index (NDVI), a widely used spectral index that quantifies vegetation “greenness” through the amount of chlorophyll, was calculated according to Rouse et al. (1974):

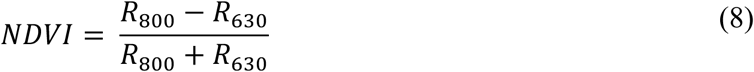

where R_800_ and R_630_ indicates reflectance at 800 and 630 nm, respectively. High NDVI values indicate high chlorophyll content, characteristic of healthy, photosynthetically active foliage (Gamon *et al*., 1995). The water index (WI) is a spectral index derived from near-infrared reflectance bands 900nm and 970nm, it was calculated according to Penuelas et al. (1997):

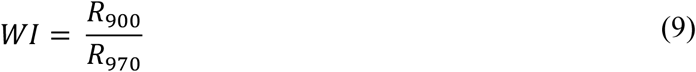

High WI values indicate greater leaf water content, as tissues with more water absorb less in the 900 and 970 nm bands (Penuelas *et al*., 1997). Consequently, elevated WI values generally reflect well-hydrated, healthy vegetation. Five needles were measured and averaged per seedling. A different portable spectrophotometer (Polypen RP410 UVIS, Photon Systems Instruments) was used for the last campaign due to technical problems with the UniSpec. Due to the limited spectral range (400-790nm) of the Polypen, WI was not calculated for the last campaign on days 98-101.

### Photosynthetic Gas Exchange Measurements

Photosynthetic gas exchange was measured on sunlit, current-year needles using a portable photosynthesis system with the opaque conifer chamber (Li-cor 6400XT and 6400-22L, Li-cor Biosciences). The system was set to a flow rate of 400 mL/min and a CO_2_ concentration of 400 ppm. Branches were pre-darkened for 30 minutes and placed into the conifer chamber with the light source set to 0 μmol quanta m^-2^ s^-1^ to quantify dark respiration (R_D_) once stability was reached at ∼5 min. Branches were then exposed to a saturating light intensity of 1500 μmol photons m^−2^ s^−1^ under 45–65% relative humidity (RH). RH was maintained at ∼60% at the ambient measurement temperature, but decreased under +5°C measurement temperature despite the use of an inline humidifier. Ambient temperature was selected based on daily averages (Table S1). Net CO_2_ assimilation (A_net_) was measured at 7-, 9-, and 11-min post-saturating light exposure and the maximum photosynthetic rate was used. Stability of gas-exchange parameters was confirmed prior to logging measurements. Recorded variables included R_D_, A_net_, stomatal conductance (g_s_), and evapotranspiration (E). Intrinsic water use efficiency (iWUE) was calculated as the ratio of A_net_ and g_s_. Immediately following measurements, needles were detached and measured for surface area using the Winseedle software package (Regent Instruments Inc.).

### Water Potential Measurements

Lower branches were excised from seedlings and kept in sealed plastic bags prior to measurement in the field. Within 3-4 minutes of sampling, pre-dawn water potential was measured on two branches per seedling using the PMS Model 1505D Pressure Chamber Instrument (PMS Instrument Company).

### Analysis of Photosynthetic Pigments

Fully expanded current-year needles were collected, flash-frozen in liquid nitrogen, and stored at - 80°C until homogenization to a fine powder in liquid nitrogen. Photosynthetic pigments were analyzed according to Junker and Ensminger (2016). Briefly, pigments were extracted from 50 mg of frozen ground needle sample in 98% methanol and 2% 0.5M ammonium acetate. Pigment extracts were analyzed using a high-performance liquid chromatography (HPLC) system (Agilent 1260 system, Agilent Technologies) with a quaternary pump, autosampler (4°C), column oven (25°C) and photodiode array detector. Pigment separation was performed on a reverse-phase C30 column (YMC Carotenoid; YMC America, Inc.) with detection at 450 and 656 nm. The mobile phase consisted of a gradient of methanol, methyl-tert-butyl-ether, and water buffered with 0.2% ammonium acetate at a flow rate of 1 ml min^-1^. For calibration and peak detection, commercially available standards were obtained from Sigma Aldrich (Oakville, ON, Canada) and DHI Lab products (Hørsholm, Denmark). Peak detection and pigment quantification were performed using ChemStation software (Agilent Technologies). Total chlorophylls (Chl) were expressed on a fresh weight basis (μmol/g). Total carotenoids per total chlorophyll (Car/Chl) were expressed as the sum of violaxanthin (V), antheraxanthin (A), zeaxanthin (Z), neoxanthin, lutein, α-carotene, and β-carotene, divided by Chl content (mmol mol^-1^). The de-epoxidation state (DEPS) of the xanthophyll cycle was calculated according to Thayer and Björkman (1990):

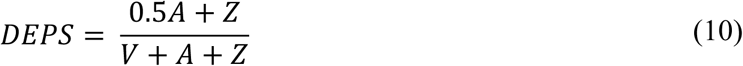

### Analysis of Xylem Development and Anatomical Traits in Early and Latewood

Xylem anatomical traits were assessed following Von Arx et al. (2016). Basal stem segments were harvested using shears during the final campaign at the end of September and stored in 80% ethanol at 4°C. Sample preparation followed a modified protocol by Chen et al. (2016; see Method S3). Cross-sections 14um thick were imaged using a Leica DVM6 digital transmitted microscope with a mid-magnification lens and LAS X v3.0.6 (Leica Biosystems) at 252x magnification for a resolution of 2.2 pixels/um. Individual images were captured with a 30% x- and y-axis overlap and a z-stack of 5 images, then stitched into a complete cross-section using PTGui Pro v11 (New House Internet Services). Two 20° xylem sectors from opposite radii were analyzed from pith to cambium using ImagePro v7 (Media Cybernetics) with the image analysis tool ROXAS v 3.0.654 (von Arx and Carrer 2014). Annual growth rings were manually delineated and assessed for primary and derived anatomical traits (see Method S4).

To assess treatment effects and family differences on xylem development within the 2021 growth ring, earlywood and latewood cells were distinguished using Mork’s Index (rTSR), defined as the radial thickness-to-span ratio of a cell, where rTSR < 1 indicates earlywood and rTSR > 1 indicates latewood (Denne, 1989). For each wood type, we quantified three xylem traits associated with hydraulic efficiency (lumen area [LA], hydraulic diameter [D_h_], theoretical hydraulic conductivity [K_h(t)_]) and three traits associated with hydraulic safety (cell wall thickness [CWT_all_], relative wall thickness [RWD], and cell wall reinforcement [(t/b)^2^]). While hydraulic efficiency and safety refer to the xylem’s capacity for water transport and resistance to cavitation, respectively (Meinzer *et al*., 2010), our measurements represent anatomical proxies rather than direct functional assessments. All analyses were conducted in R v.4.4.3 (R Core Team, 2025).

### Height and Radial Stem Growth

During each measurement campaign, height and radial stem diameter were measured for all seedlings in all plots. Height was measured from the base at the soil surface to the highest point on the main stem using a tape measure, while radial stem diameter was measured at the base of the stem near the soil surface with standard digital calipers. One seedling per family per plot was harvested the following spring and dried to a constant mass at 65°C. Seedlings were divided into roots, shoots, and leaves, and each tissue was weighed individually.

### Statistics

Three-way ANOVA models were used to assess the effect of temperature, VWC and family on all measured parameters, with plot included as a random factor for each measurement campaign. Each plot was used as a biological replicate. Linear mixed-effects ANCOVA models were used to assess the effect of temperature, VWC and family on height and diameter growth over the experiment, accounting for the effect of seasonal variation introduced by minimum ambient daily air temperature and photoperiod. The ANCOVA models used temperature, VWC, and family as categorial fixed factors, photoperiod and minimum ambient daily air temperature as continuous numeric covariates, and plot and ID as nested random factors.

All ANOVA and ANCOVA models were run using *lme4* (Bates *et al*., 2015) in R v.4.4.3 (R Core Team, 2025). Post-hoc Tukey pairwise comparisons were performed on ANOVA models (P<0.05) using estimated marginal means in the *emmeans* package (Lenth, 2023). For each parameter, the best-fit model was selected based on the lowest Akaike information criterion (AIC) using *AICcmodavg* (Mazerolle 2023). Maximum likelihood estimation was used to prioritize accurate estimates of the fixed regression parameters. Data preprocessing was conducted with *dplyr* (Wickham *et al.,* 2023) and figures were generated using *ggplot2* (Wickham *et al*., 2023).

## Results

### Environmental Conditions

Conditions in the plots the day before the rainout (RO) treatment began showed mean daily air temperature, leaf temperature, VPD, and soil moisture values of 15.9°C, 15.4°C, 1.46kPa, and 0.24 m^3^/m^3^ (Fig. **2a–d**). RO structures installed on June 24 lowered soil moisture in RO plots to 0.18 m^3^/m^3^ by August 15, while control plots were maintained at 0.26 ± 0.01 m^3^/m^3^. Infrared warming began on August 16 and had minimal impact on soil moisture. Later in August, when ambient air temperatures surpassed 30 °C, soil moisture reached its lowest levels: 0.16 m^3^/m^3^ in RO plots and 0.15 m^3^/m^3^ in RO+W plots, compared with 0.25 m^3^/m^3^ in control plots. Heavy rainfall near the end of the season (Fig. **2d**) increased soil moisture across treatments, though deficits in RO and RO+W plots persisted, averaging 0.21 m^3^/m^3^ and 0.19 m^3^/m^3^ and remaining 22% and 27.9% lower than controls across the warming period.

The warming treatment raised daily mean air temperatures (T_air_) by an average of 1.3°C, with maximum increases of 2.1°C (Fig. **2a**) Daily minimum T_air_ responded more strongly, rising by 2.4°C on average and up to 4°C. Estimated daily T_leaf_ was increased by an average of 3.9°C, with maximum increases of 4.7°C observed (Fig **2b**). In addition to the soil-moisture deficit described above, warming raised vapour pressure deficit (VPD), on average from 1.71 kPa in control plots to 1.77 kPa in warming plots (Fig**. 2c**).

### Growth and Biomass Accumulation

The fast-growing family exhibited greater height and diameter growth throughout the experiment compared to the slow-growing family (Fig. **3a,b**; Table S2). Aboveground biomass was also higher in the fast-growing family, while belowground biomass and total biomass did not differ significantly between families (Fig. **3c,d****,e**; Table S2). The root: shoot ratio was higher in the control treatment of the slow-growing family compared to the fast-growing family, but this difference was not statistically significant (Fig**. 3f**; Table S2). RO, warming, and their combination had negligible effects on height, diameter growth and biomass accumulation in either family.

**Figure 3.**
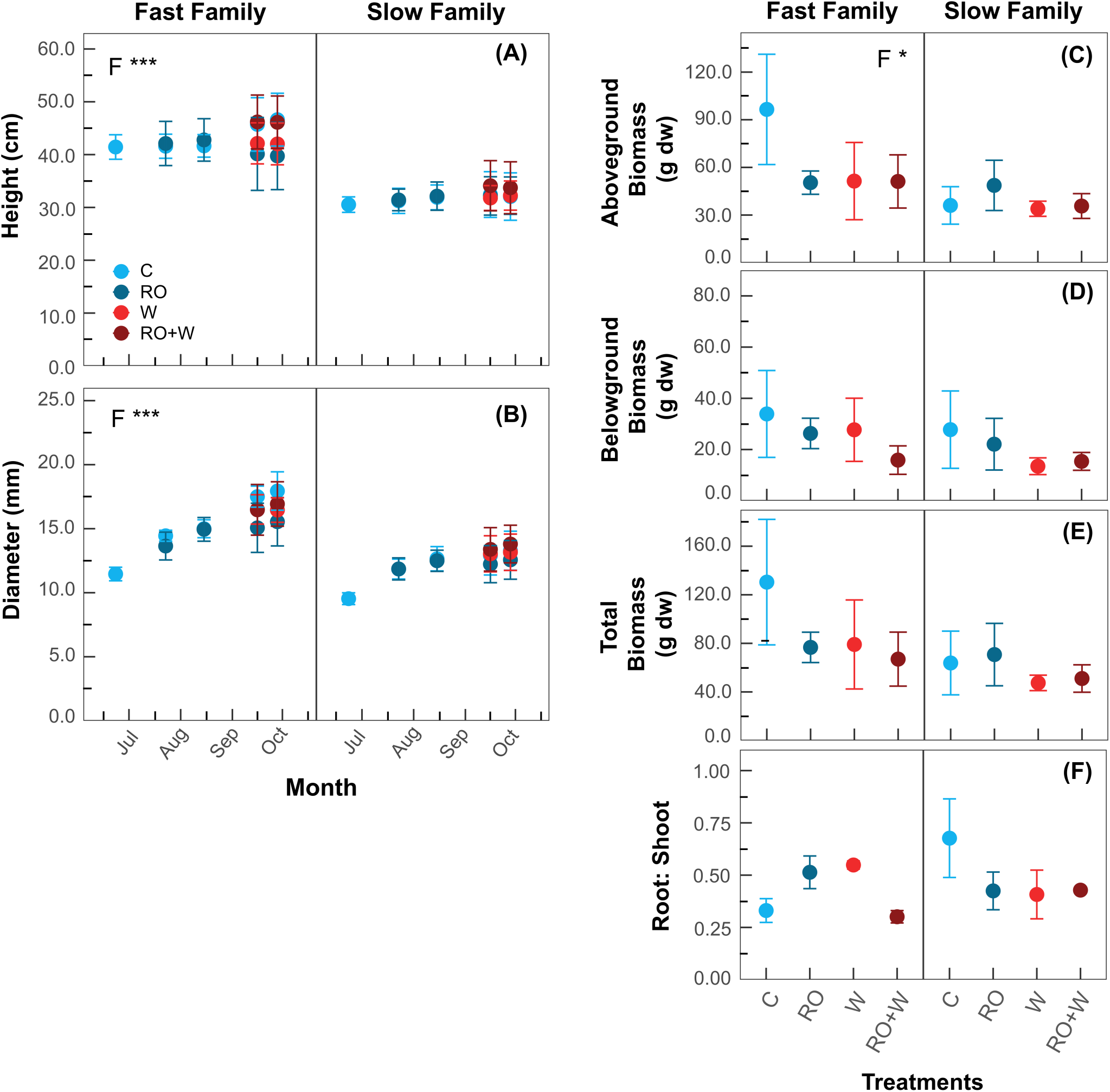
The impacts of summer rainout and warming treatments on growth in a fast- and slow-growing family of white spruce. Control, C; Rainout, RO; Warming, W; Rainout combined with warming, RO+W. (**a**) height at each measurement date; (**b)** diameter at each measurement date; (**c**) final aboveground biomass (g dw); **(d)** final belowground biomass (g dw); **(e**) final total biomass (g dw); and (**f**) the ratio of root to shoot biomass at the end of the experiment. Data points represent the average of three plot; n=3, ± SE. Temperature (T), volumetric water content (VWC), and family (F) were tested as main effects. Only significant main effects or interactions are noted (P<0.05). *P < 0.05, **P < 0.01 and ***P < 0.001.

### Xylem Anatomical Traits in Early- versus Latewood

Earlywood xylem traits were consistent across families and were unaffected by combined RO and warming at the end of the experiment. Anatomical proxies for hydraulic efficiency, including lumen area (LA), hydraulic diameter (D_h_), and theoretical hydraulic conductivity (K_h(t)_), and for hydraulic safety, including cell wall thickness (CWT) and tracheid wall reinforcement ([t/b]^2^) did not differ between the fast- and slow-growing families (Fig. **4a,c****,e,g,i;** Table S2). Mechanical support, estimated via relative wood density (RWD), also showed no family-level differences in earlywood (Fig. **4k**).

**Figure 4.**
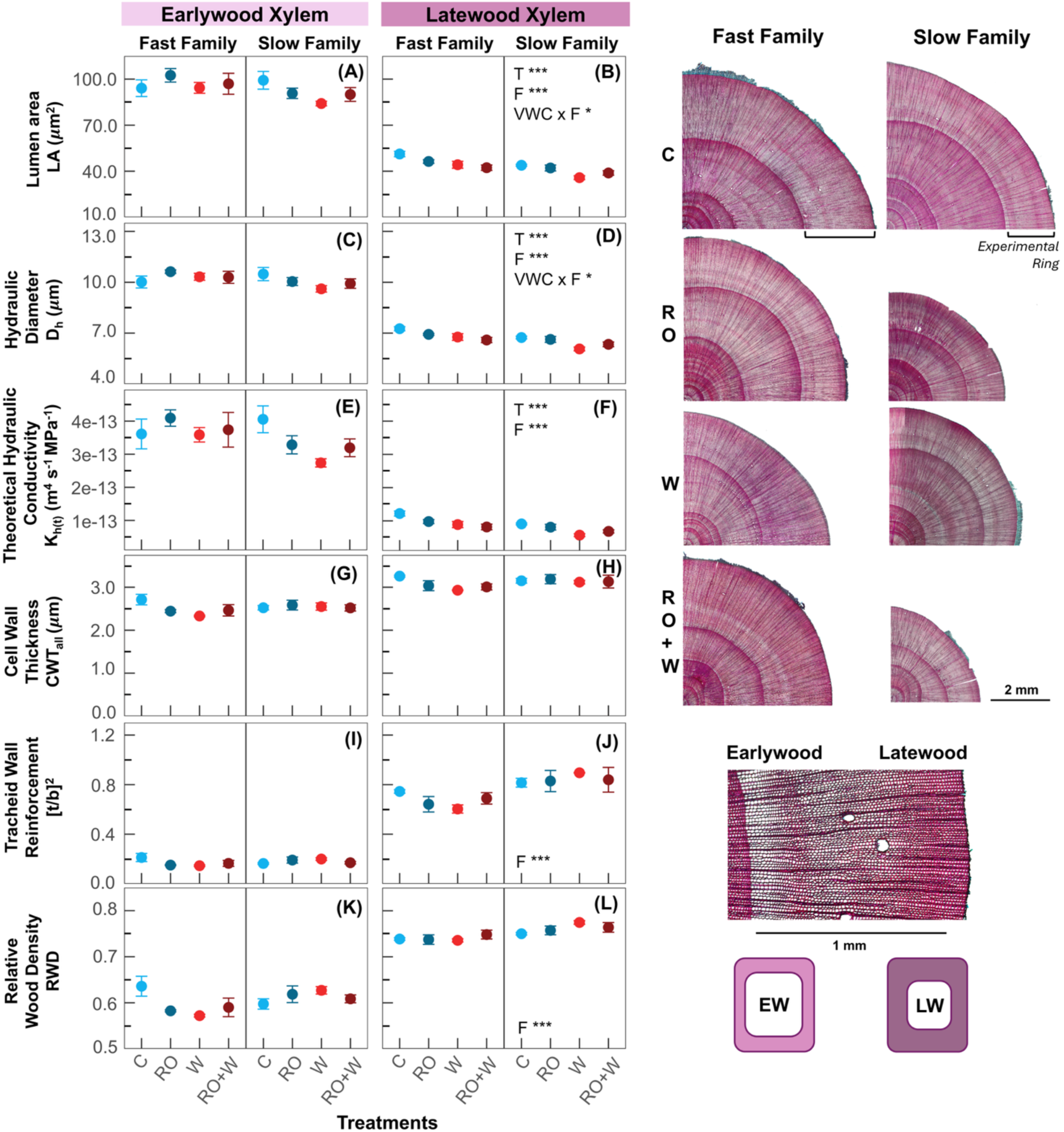
The effects of summer rainout and warming treatments on xylem earlywood and latewood development in a fast-growing and slow-growing white spruce family. Hydraulic conductivity, characterized by **(a,b)** lumen area (LA), **(c,d)** hydraulic diameter (D_h_), and (**e,f**) theoretical specific conductivity (K_h(t)_). Hydraulic safety, characterized by **(g,h)** cell wall thickness (CWT_all_); **(i,j)** tracheid wall reinforcement [t/b]^2^. Mechanical support, characterized by: **(k,l)** relative wood density (RWD). Control, C; Rainout, RO; Warming, W; Rainout combined with warming, RO+W. Measurements were averaged from two 20° sectors from opposite radii of the 2021 growth ring. Data points represents the average of 3 plots; n=3, ±SE. Temperature (T), volumetric water content (VWC), and family (F) were tested as main effects. Only significant main effects or interactions are noted (P<0.05). *P < 0.05, **P < 0.01 and ***P < 0.001. K_h(t)_ values were log-transformed prior to statistical analysis to improve numerical stability and interpretability due to their very small magnitude.

In contrast to the earlywood, latewood xylem traits revealed significant family-level differences and responses to individual stress treatments. Overall, the fast-growing family had higher LA, D_h_, and K_h(t)_ but lower [t/b]^2^ and RWD compared to the slow-growing family (Fig. **4b,d,f,j,l**; Table S2). Warming reduced latewood LA, D_h_ and K_h(t)_ across both families, while RO had a family specific-effect, with the fast-growing family exhibiting greater reductions in LA and D_h_ than the slow-growing family. Anatomical proxies for hydraulic efficiency and safety remained unchanged under the combined treatment.

### Pre-dawn Water Potential

Pre-dawn water potential was similar in both families when the experiment started and throughout the experiment (Fig. **5**; Table S3). Warming significantly reduced pre-dawn water potential by ∼33% in both families during the last measurement campaign (Fig. **5c**). However, RO had no measurable impacts on water potential during the experiment, despite a prolonged reduction in soil moisture (Fig. **2d**). Overall, water potential remained higher than −0.5 MPa across all treatments and time points.

**Figure 5.**
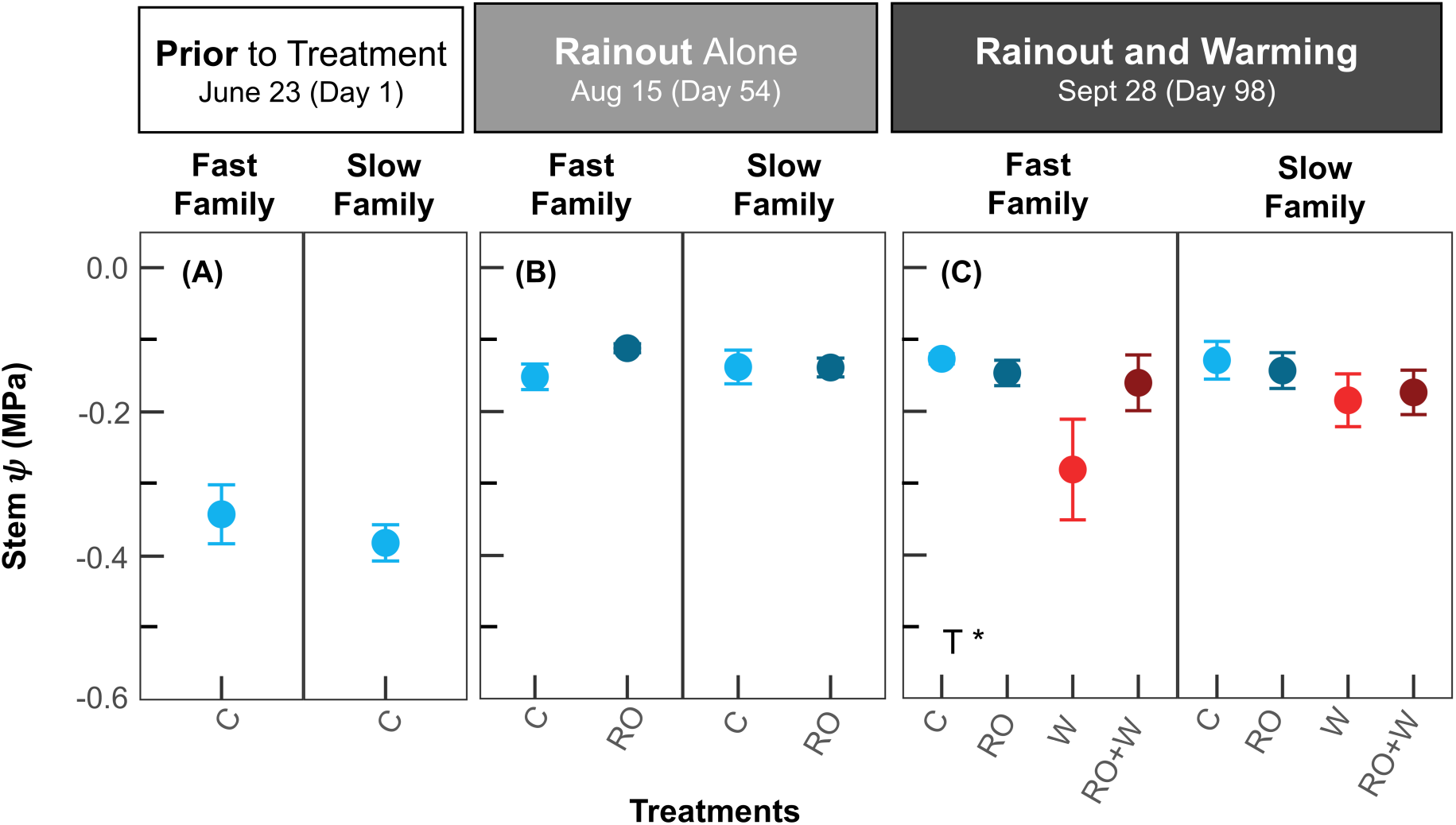
Effects of summer rainout and warming treatments on pre-dawn stem water potential in a fast-growing and slow-growing family of white spruce. Data points represent the average of 12 plots for Jun 23, 5-6 plots for Aug 15, and 2-3 plots for Sept 28, ±SE. Control, C; Rainout, RO; Warming, W; Rainout combined with warming, RO+W. Temperature (T), volumetric water content (VWC), and family (F) were tested as main effects with minimum temperature as a covariate. Only significant main effects or interactions are noted (P<0.05). *P < 0.05, **P < 0.01 and ***P < 0.001.

### Photosynthetic Gas Exchange

Despite differences in growth strategies, both families had similar net photosynthetic rates (A_net_), dark respiration (R_D_), and stomatal conductance (g_s_) prior to treatment initiation and through most of the experiment (Fig. **6a-l**; Table S3). Family-level differences were only briefly apparent in August, when the fast-growing family had higher g_s_ and transpiration (E) compared to the slow-growing family (Fig. **6k,o**; Table S3). The fast-growing family also exhibited a transient reduction in intrinsic water use efficiency (iWUE) under RO compared to the slow-growing family, which disappeared after seven weeks (Fig. **6r**; Table S3). By September, all family differences had diminished and both families responded similarly to combined warming with RO.

**Figure 6.**
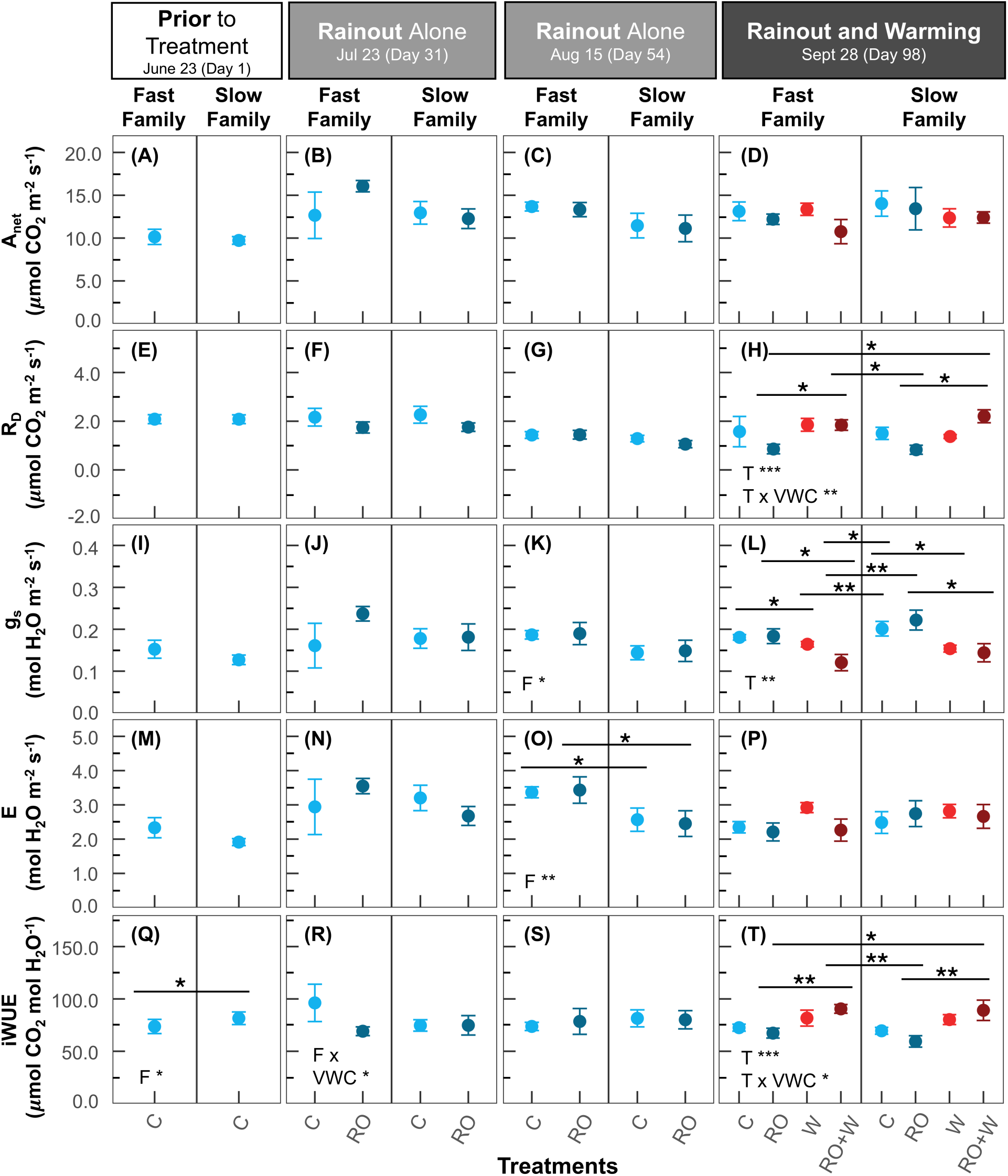
Effects of summer rainout and warming treatments on photosynthetic gas exchange in a fast-growing and slow-growing family of white spruce. **(a-d)** Net photosynthetic rate (A_net_); **(e-h)** dark respiration rate (R_dark_); **(i-l)** stomatal conductance (g_s_); (**m-p**) transpiration rate (E); (**q-t**) intrinsic water use efficiency (iWUE). Data points represent the average of 12 plots for Jun 23, 5-6 plots for July 23 and Aug 15, and 2-3 plots for Sept 28, ±SE. Control, C; Rainout, RO; Warming, W; Rainout combined with warming, RO+W. Temperature (T), volumetric water content (VWC) and family (F) were tested as main effects. Only significant main effects or interactions are noted (P<0.05). *P < 0.05, **P < 0.01 and ***P < 0.001.

Seedlings exposed to and measured under 5°C warming maintained similar net photosynthetic rates (A_net_) comparable to control seedlings, with no significant impact observed under RO (Fig**. 6d**; Table S3). In contrast, R_D_ rates increased under warming, with the highest rates observed under the combined warming and RO treatment (Fig. **6h**; Table S3). Warming also led to a significant reduction in g_s_ (Fig **6l**; Table S3) accompanied by a significant increase in iWUE with the highest values observed under warming and RO (Fig. **6t**; Table S3). RO alone had minimal effects on leaf-level gas exchange.

### Chlorophyll Fluorescence

Throughout most of the summer, chlorophyll fluorescence parameters remained similar between families (Fig. **7**; Table S3). By late September, however, the slow-growing family had higher yield of photosystem II (*φ*_PSII_), lower dynamic non-photochemical quenching (*φ*_NPQ_) and excitation pressure on PSII (1-qP) compared to the fast-growing family (Fig. **7h,l,t**; Table S3). Family-specific responses to RO also emerged in sustained non-photochemical quenching (*φ*_f,D_) and 1-qP, with both parameters being higher in the slow-growing family and lower in the fast-growing family under RO conditions (Fig. **7p,t**; Table S3). Small but significant effects of only warming or only RO were also observed in several non-photochemical quenching parameters (Fig. **7**; Table S3). F_v_/F_m_ and *φ*_PSII_ were both elevated in the RO treatment compared to the control, 54 days after the RO treatment began (Fig. **7c,g**; Table S3). However, values of F_v_/F_m_ and *φ*_PSII_ remained relatively high across treatments, exceeding 0.77 and 0.22 respectively. By the end of the experiment, sustained NPQ was significantly higher under warming in the slow-growing family, specifically between the C and RO+W groups (P=0.011), though values remained below 0.25 (Fig. **7p**; Table S3).

**Figure 7.**
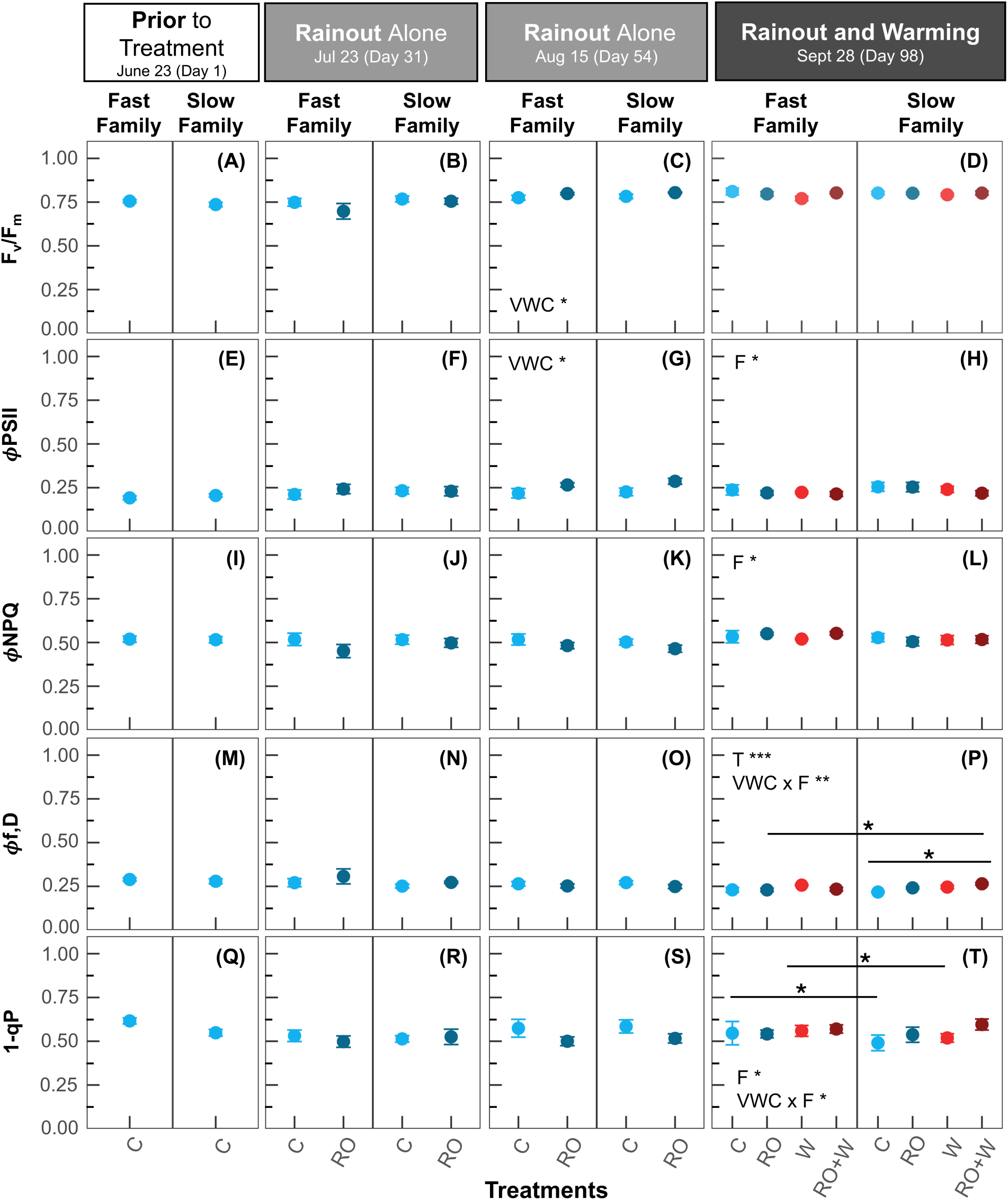
Effects of summer rainout and warming treatments on chlorophyll fluorescence in a fast-growing and slow-growing family of white spruce. **(a-d)** Maximal quantum yield of PSII (F_v_/F_m_); (**e-h**) effective quantum yield of PSII (ϕ_PSII_); (**i-l**) dynamic non-photochemical quenching (ϕ_NPQ_); (**m-p**) the sum of fluorescence and sustained non-photochemical quenching (ϕ_f,D_); (**q-t**) excitation pressure of the reaction centers of PSII (1-qP). Data points represent the average of 12 plots for Jun 23, 6 plots for July 23 and Aug 15, and 3 plots for Sept 28, ±SE. Control, C; Rainout, RO; Warming, W; Rainout combined with warming, RO+W. Temperature (T), volumetric water content (VWC) and family (F) were tested as main effects. Only significant main effects or interactions are noted (P<0.05). *P <0.05, **P < 0.01 and ***P < 0.001.

### Spectral Reflectance

For most of the experiment, we did not observe treatment effects on the four measured vegetation indices; however, an interactive effect of family and RO on spectral reflectance did emerge at the end. Prior to the RO treatment initiation in June, the fast- and slow-growing white spruce families exhibited similar values for photochemical reflectance index (PRI), chlorophyll carotenoid index (CCI) and normalized difference vegetation index (NDVI; Fig. **8a,e,i**; Table S3). From July to mid-August, PRI was significantly higher in the fast-growing family, while CCI and NDVI remained similar between families (Fig. **8b,c**; Table S3). The water index (WI) was consistently higher in the fast-growing family across all three time points, indicative of higher water content in the needles, relative to the slow-growing family (Fig. **8m-o**; Table S3). In September, following six weeks of RO combined with warming, PRI, CCI, and NDVI were higher in the fast-growing family and lower in the slow-growing family under the soil-moisture deficit (Fig. **8d,h,l**; Table S3). Notably, none of the pairwise comparisons were statistically significant.

**Figure 8.**
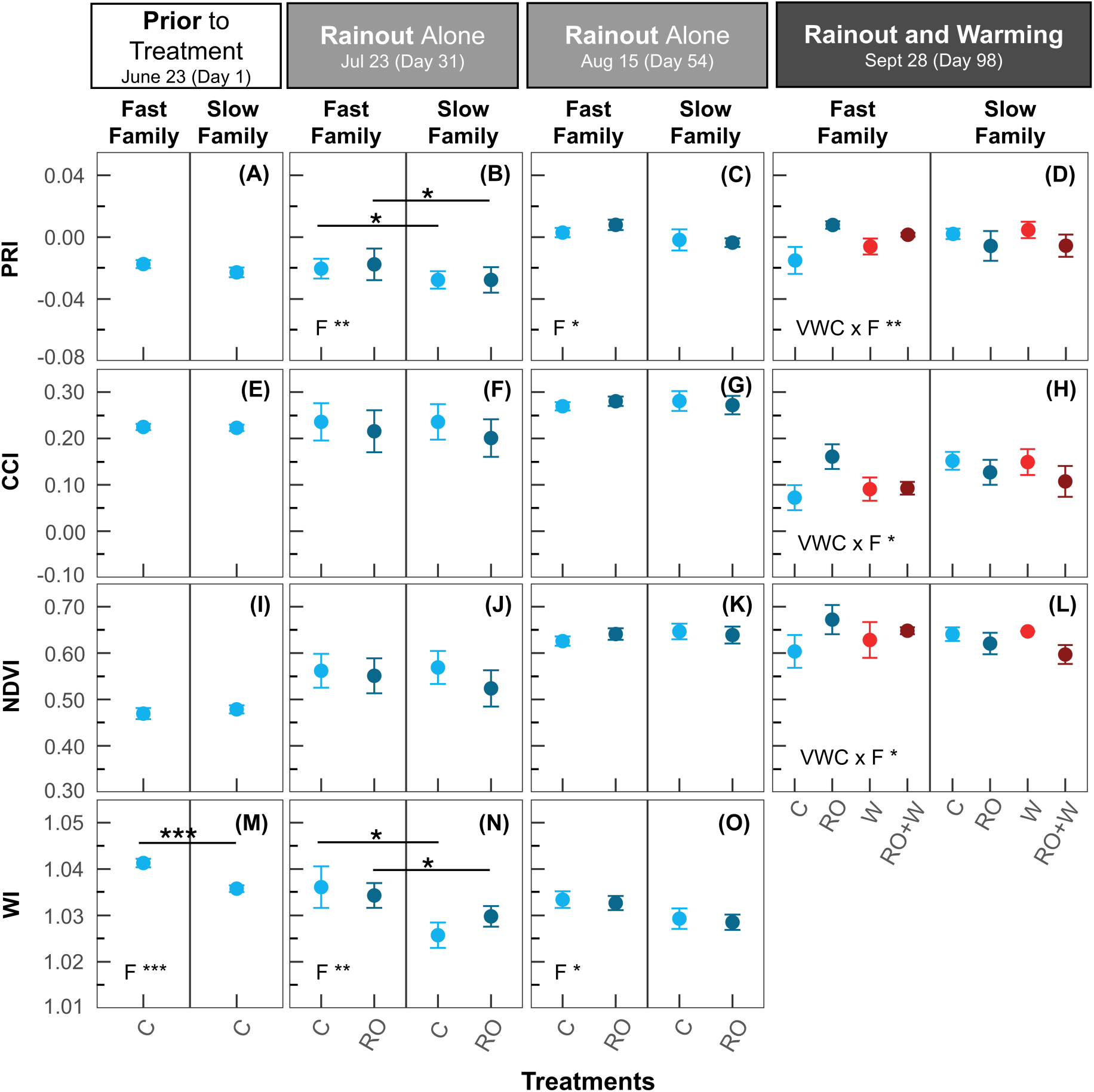
Effects of summer rainout and warming treatments on photosynthesis as observed through spectral reflectance indices in a fast-growing and slow-growing family of white spruce. **(a-d)** Photochemical reflectance index (PRI); (**e-h**) Chlorophyll carotenoid index (CCI); (**i-l**) normalized vegetation index (NDVI); (**m-o**) water index (WI). Data points represent the average of 12 plots for Jun 23, 6 plots for July 23 and Aug 15, and 3 plots for Sept 28, ±SE. Control, C; Rainout, RO; Warming, W; Rainout combined with warming, RO+W. Temperature (T), volumetric water content (VWC) and family (F) were tested as main effects. Only significant main effects or interactions are noted (P<0.05). *P < 0.05, **P < 0.01 and ***P < 0.001.

### Photosynthetic Pigments

Family-level differences in pigment composition were evident at both sampling times. In August, the fast-growing family showed a pronounced increase in chlorophyll under RO (Fig 9**a**; Table S3), and higher overall lutein concentrations, lower zeaxanthin concentrations, and a lower de-epoxidation state compared to the slow-growing family (Fig. **9e,g,k**; Table S3). By September, total chlorophyll remained elevated in the fast-growing family, while total carotenoids and xanthophyll pigments were lower relative to the slow-growing family (Fig. **9b,d,j**; Table S3).

**Figure 9.**
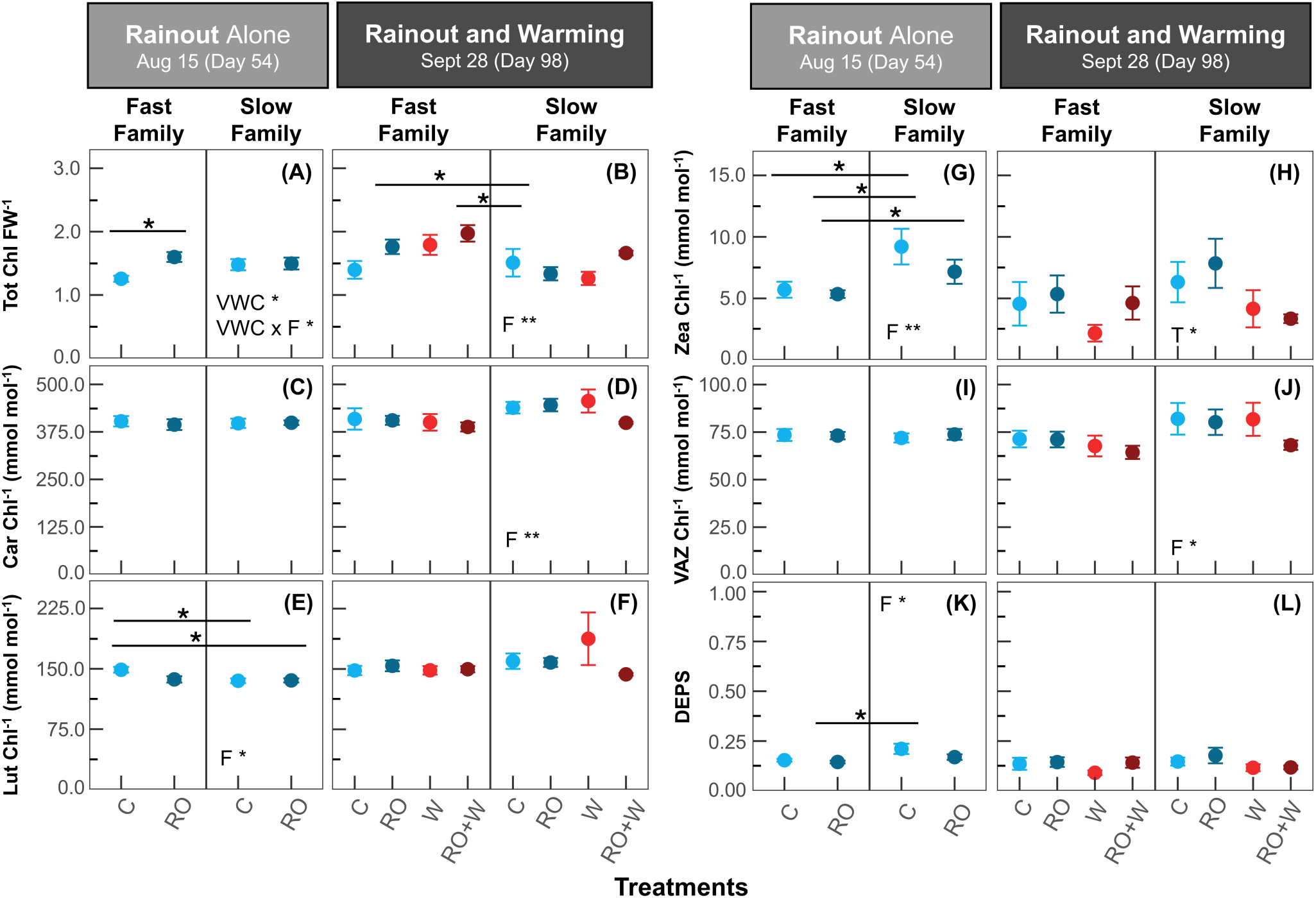
Effects of summer rainout and warming treatments on photosynthetic pigments in a fast-growing and slow-growing family of white spruce. **(a-b**) Total chlorophyll per fresh weight (Chl FW^-1^); (**c-d**) ratio of carotenoids to chlorophylls (Car Chl^-1^); (**e-f**) ratio of lutein to chlorophylls (Lut Chl^-1^); (**g-h**) ratio of zeaxanthin to chlorophylls (Zea Chl^-1^); (**i-j**) ratio of all xanophyll cycle pigments to chlorophylls (VAZ Chl^-1^); (**k-l**) de-epoxidation state (DEPS). Data points represent the average of 6 plots for Aug 15 and 3 plots for Sept 28, ±SE. Control, C; Rainout, RO; Warming, W; Rainout combined with warming, RO+W Temperature (T), volumetric water content (VWC), and family (F) were tested as main effects with minimum temperature as a covariate. Only significant main effects or interactions are noted (P<0.05). *P < 0.05, **P < 0.01 and ***P < 0.001.

The effect of combined warming and RO treatment on pigment composition in white spruce seedlings was small. In August, total chlorophyll concentration was higher in the RO seedlings compared to the control seedlings (Fig. **9a**; Table S3), but this effect was no longer evident by September. Carotenoid and xanthophyll cycle pigment concentrations were unaffected by RO at either time points (Fig. **9c-j**; Table S3). Warming alone led to lower amounts of zeaxanthin in September, but had only little impact on other photosynthetic pigments (Fig. **9h**; Table S3).

## Discussion

### Fast-growing family retains growth advantage under concurrent warming and soil-moisture deficit

To what extent does genetic variation within a species influence resilience to combined climatic stress? Although species-level responses among conifers are well-studied, little is known about how intraspecific variation, particularly in white spruce, shapes resilience to concurrent warming and reduced soil moisture. The broad geographic ranges of boreal conifers expose populations to diverse environmental conditions, which drive local adaptation and generate trait variation linked to a greater tolerance to heat and drought. In our experiment, the fast-growing white spruce family retained a substantial growth advantage, exhibiting on average 75% greater height growth and 42% greater diameter growth, resulting in higher aboveground biomass (Figs. **S3** and **3**). These growth differences were consistent across treatments. However, contrary to our first hypothesis, both families had comparable rates of photosynthesis, stomatal conductance, and water potential under warming, rainout, and their combination, revealing a surprising resilience to the simulated climatic stress irrespective of growth strategy.

Conifers adapt to diverse environments by modifying xylem morphology (Pittermann *et al*., 2012; Bouche *et al*., 2014; Zheng *et al*., 2022), prompting our investigation into xylem traits related to hydraulic safety and efficiency. Consistent with this adaptive capacity demonstrated previously at the species level, we observed family-level differences in xylem anatomy specifically related to hydraulic efficiency. A widely debated hypothesis in plant hydraulics posits a functional trade-off between hydraulic safety and efficiency (e.g., Hacke and Sperry, 2001; Gleason *et al*., 2012; Venturas *et al.,* 2017; Prendin *et al*., 2018). According to this framework, xylem traits that reduce the risk of drought-induced hydraulic failure are often associated with lower hydraulic efficiency, which constrains growth. Although this framework is typically evaluated at the interspecific level, our findings demonstrate that similar trade-offs can be observed at the intraspecific level within white spruce. The fast-growing family had higher values for xylem traits related to hydraulic efficiency, such as lumen area (LA), hydraulic diameter (D_h_), and theoretical hydraulic conductivity (K_h(t)_) and lower values for xylem traits related to hydraulic safety, including tracheid wall reinforcement and relative wood density (Fig. **4b,d,f,j,l**). These family-level differences were confined to latewood, which form later in the growing season and is under moderate genetic control in white spruce (Soro *et al*., 2022).

Despite anatomical patterns consistent with a hydraulic safety-efficiency trade-off (Fig **S4**), both families maintained hydraulic function under concurrent warming and reduced soil moisture, as indicated by high water potentials (Fig. **5**). Notably, traits related to hydraulic efficiency in the latewood varied by family: the slow-growing family increased latewood D_h_ and LA under the rainout treatment, whereas the fast-growing family showed a reduction in these traits (Fig. **4b,d**). This pattern suggests that the ability to maintain hydraulic function may rely more on plasticity in traits associated with hydraulic efficiency, rather than direct shifts in traits linked to hydraulic safety (e.g., cell wall thickness). Wood formation in mature silver fir and Scots pine has shown to be strongly constrained by water availability during hot years (Larysch *et al*., 2022). Here, we demonstrate that white spruce exhibits intraspecific variation in these anatomical traits, and that such variation is evident even under more moderate climatic stress. Plasticity in wood formation is widespread among conifers and is often expressed through the development of latewood tracheids with smaller lumen and thicker cell walls, yielding denser wood during drought years (Huang *et al*., 2022). In our study, warming also led to reductions in LA, K_h(t),_ and D_h_ in white spruce seedlings (Fig **4b,d,f**). During hot, dry summers, reduced turgor pressure can limit cambial expansion, resulting in smaller-diameter tracheids with thicker cell walls, a response that contributes to maintaining water transport under stress (Huang *et al*., 2022). Interestingly, resilience to concurrent soil moisture deficits and warming in white spruce seedlings in our experiment was achieved through only small anatomical adjustments in anatomical traits related to hydraulic efficiency.

Beyond xylem anatomy, additional family-level differences in response to soil-moisture deficit and warming emerged in leaf-level gas exchange and spectral reflectance during mid- to late summer, with reflectance-derived vegetation indices being the most sensitive for revealing genetic variation. The fast-growing family consistently had higher values in the photochemical reflectance index (PRI) and water index (WI) from July to August (Fig. **8b,c,m-o**). Higher PRI typically reflects lower engagement of the photoprotective xanthophyll cycle and reduced conversion of violaxanthin to zeaxanthin, indicating less non-photochemical quenching (NPQ) and therefore lower stress (Fréchette *et al*., 2016, 2020; D’Odorico *et al*., 2021) This pattern is consistent with our pigment data, as the fast-growing family exhibited lower zeaxanthin levels when PRI was elevated (Fig. **9g**). Taken together, the higher PRI in the fast-growing family suggests that its photosynthetic apparatus operated more efficiently and experienced less excess light energy than that of the slow-growing family. Interestingly, NPQ did not differ between families, likely reflecting a decoupling between pigment-level variation captured by PRI and mechanisms of NPQ regulation not fully detected by PRI (Porcar-Castell *et al*., 2012; Fréchette *et al*., 2015). By the end of the growing season, the chlorophyll carotenoid index (CCI), normalized difference vegetation index (NDVI), and PRI all showed intraspecific responses to the rainout treatment, with the fast-growing family maintaining higher values whereas values declined in the slow-growing family. Together, these findings demonstrate that spectral vegetation indices reliably trace variation in leaf traits, with PRI, WI, and pigment dynamics converging to capture consistent family-level differences in white spruce under soil-moisture deficits.

### A warmer, drier climate is within the adaptive capacity of white spruce

Contrary to our second hypothesis that concurrent warming and soil-moisture deficits would most strongly suppress photosynthesis and growth, five-year-old white spruce seedlings were highly resilient to warming and rainout, individually and in combination. Photosynthetic rates, photochemical efficiency, water potential, and growth remained consistently high across treatments (Figs. **3****, 5-7**). Given the realistic, moderate intensity of our applied treatments, these findings contribute to emerging evidence that heat and drought synergistically intensify tree stress only once conditions exceed species’ hydraulic or metabolic limits. In interspecific analyses of evergreen and deciduous northern taxa, only extreme warming (+7.7°C) coupled with repeated or extended drought reduced first-year seedling survival, with no growth declines among survivors (MacDonald *et al*., 2026). In pine seedlings, gradually intensifying extreme heat (30°C to 40 °C) combined with drought more strongly and reduced photosynthesis, photochemical efficiency, and water potential than under high heat alone (Rehschuh & Ruehr, 2022). This raises a key question: were conditions simply below stress thresholds or did phenotypic plasticity preserve seedling performance?

In our field experiment, a ∼50% reduction in rainfall did not impose sufficient soil-moisture limitations to exceed the drought tolerance of white spruce seedlings, and therefore the combined warming and rainout treatment also failed to elicit strong stress responses. Although rainout structures effectively reduced soil moisture (Fig. **2d**), pre-dawn stem water potential remained above –0.5 MPa across treatments (Fig. **5**), indicating adequate hydration even under the combination of rainout and warming. White spruce seedlings exhibited adjustments consistent with drought acclimation that likely helped them to maintain hydraulic and photosynthetic function. After seven weeks of progressive soil drying, rainout seedlings had higher total chlorophyll and elevated F_v_/F_m_ and Φ_PSII_ relative to controls (Figs. **7c,g** and **9a**). With net CO_2_ assimilation unchanged (Fig. **6d**), increased chlorophyll likely buffered photosynthetic capacity under drier conditions. Given that the applied warming and precipitation reductions align with projected summer conditions in much of eastern Canada (Wang *et al*., 2022), these responses point to ecologically relevant plasticity and resilience in white spruce.

Several conifer species, including Norway spruce, Silver fir, Douglas fir and Altas cedar, have been classified as drought-tolerant but heat-sensitive based on thermal limits of F_v_/F_m_ and leaf turgor loss point (Kunert *et al*., 2022). With sufficient water, these species can partially offset thermal risk by sustaining evaporative cooling during heatwaves, keeping leaf temperatures below damaging levels. Even so, this heat sensitivity is relative. For example, Norway spruce exhibited the lowest temperature threshold for the onset of F_v_/F_m_ decline at 38.5±0.8°C. In our field study, air temperatures did not reach this threshold in control or heated plots (Fig. **2a**). Nevertheless, warming alone and combined with reduced soil moisture had pronounced effects on leaf gas exchange. An applied +5°C increase in leaf temperature significantly reduced stomatal conductance (g_s_), yet seedlings exposed to warming maintained similar photosynthesis (A_net_) and thus achieved higher intrinsic water use efficiency than those under ambient conditions (Fig. **6d,l,t**). Similarly, under severe drought, most mature white spruce families are able to maintain or increase iWUE via regulation of g_s_ (Depardieu *et al*., 2024). These g_s_-driven gains in iWUE raise a mechanistic question: is maintained A_net_ explained solely by reduced g_s_ or do adjustments in photosynthetic capacity also contribute?

Carbon uptake can be maintained under warming by increasing the thermal optimum of photosynthesis (T_optA_; Berry and Bjorkman 1980; Gunderson *et al.,* 2010; Way and Yamori 2014; Sendall *et al.,* 2015). Such acclimation mainly reflects shifts in the temperatures at which carboxylation and electron transport operate most efficiently, rather than by changes in stomatal conductance or respiration alone (Kumarathunge *et al*., 2019; Dusenge *et al*., 2023). Although our observations point to increased iWUE via lower g_s_, distinguishing stomatal from capacity-driven acclimation in white spruce requires direct measurements of T_optA_, the maximum carboxylation capacity of Rubisco (V_cmax_), and electron transport capacity (J_max_) in future studies.

Unlike short-term heatwaves in controlled studies, typically four to 14 days with temperature increases around 10°C (Ameye *et al.,* 2012; Duarte *et al.,* 2016; Birami *et al.,* 2018; Guha *et al.,* 2018), our six-week warming treatment provided a prolonged and ecologically realistic thermal regime that likely permitted gradual thermal acclimation. In a greenhouse experiment on white spruce, Gagné *et al*. (2020) observed reduced gas exchange under combined heat and drought with effects diminishing over repeated exposure. This attenuation of stress mirrors our findings and suggests that extended or prior exposure facilitates the acclimatory adjustments needed to sustain photosynthetic function and tolerance to concurrent stress. Although both studies observed elevated respiration under combined warming and soil-moisture deficits (Fig. **6h**), the more extreme +10 °C warming in Gagné *et al*. likely amplified carbon losses and reduced biomass, outcomes we did not observe under our more moderate +5°C warming.

### Conclusions

Simulating future summer conditions projected for southeastern Canada, we found that both fast- and slow-growing white spruce families were resilient under soil moisture as low as 15% and air temperatures up to 34.5°C. Our results reveal that coordinated adjustments in xylem architecture, pigment pools, and photosynthetic traits allowed seedlings to sustain gas exchange and growth during prolonged warming and soil-moisture deficit (Fig. **10**). Higher intrinsic water-use efficiency from reduced stomatal conductance, together with smaller latewood xylem lumen areas, contributed to this coordinated response. The contrast in growth strategy (fast vs. slow) explained more variance in growth and xylem development than treatment effects, indicating that faster growth does not increase vulnerability in white spruce at mild to moderate stress levels. Growth rate alone may therefore be a poor indicator of resilience to warming and drought when selecting families for future reforestation, given the high plasticity observed at the seedling stage.

**Figure 10.**
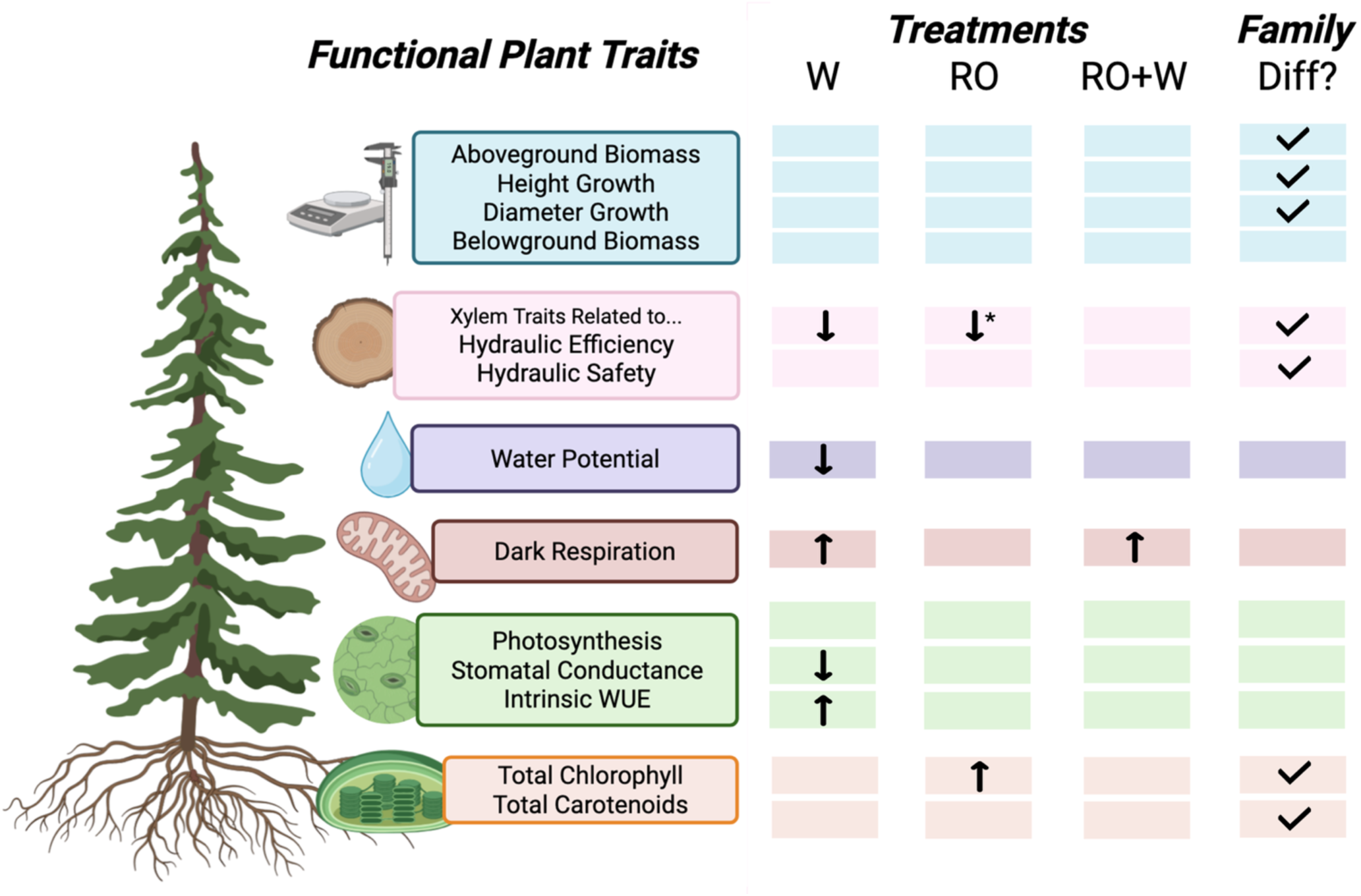
Schematic summary of treatment effects and family differences on functional plant traits under warming (W), rainout (RO), and their combination (RO+W). Arrows indicate trait responses, and arrows labeled with “*” denote significant treatment × family interactions, highlighting family-specific responses. WUE, water use efficiency. Created in BioRender. Murphy, B. (2026) https://BioRender.com/ettlls2.

## Acknowledgements

We are grateful to the support provided by the Canadian Forest Service Quebec, in particular to Daniel Plourde and Eric Dussault for seedling production, planting, and technical support during setup of the experiment. Authors are also grateful to Georg von Arx for his assistance with ROXAS; Maia Dall-Acqua and Anchalya Balasubramaniam for help in the field; Helena Jovic for help processing xylem samples; Phil Rudz and Mirek Szreder for technical support with the T-FACE set-up; and Kate Brown for maintenance support at the field site.

## Author Contributions

I.E., N.I. and B.K.M. conceptualized the experiment. B.K.M., F.N., N.P., S.W., T.M., and J.A.B performed all experimental work. B.K.M. analyzed the data. B.K.M. and I.E. wrote the manuscript. All authors have reviewed the manuscript.

## Data Availability

The data underlying this article will be shared on reasonable request to the corresponding author.

## Supporting Information

**Methods S1**. Installed sensors in KSR plots

**Methods S2.** Regression-based estimation of leaf temperature

**Methods S3.** Xylem sample preparation

**Methods S4.** Primary xylem and derived anatomical traits assessed by ROXAS

**Fig. S1.** Seasonal temperatures and annual precipitation for the period 2000-2020 at the parental origin sites for the two families of white spruce.

**Fig. S2.** The temperature free-air-enhancement and rainout experimental set up at the Koffler Scientific Reserve in Ontario, Canada.

**Fig. S3.** The impacts of summer rainout and warming on aboveground growth in a fast- and slow-growing genotype of white spruce.

**Fig. S4.** Correlations between primary and derived xylem anatomical traits related to hydraulic efficiency and hydraulic safety in a fast- and slow-growing family of white spruce.

**Fig. S5.** Approximate points where manual soil volumetric water content (VWC) was measured and then averaged from in each plot.

**Fig. S6.** Automatic soil sensor corrections applied for the 2021 experimental time frame from June 24 to October 2 at the Koffler Scientific Reserve in Ontario, Canada.

**Fig. S7.** Observed versus predicted withheld values from a single 50/50 train–test validation of regression models developed using 2020 data from Murphy *et al. (*2025).

**Fig. S8.** Comparison of measured and estimated air and leaf temperatures across experiments.

**Table S1.** Cuvette temperature and relative humidity measured during each measurement campaign using the LI-6400 XT gas exchange system.

**Table S2.** Summary of Three-Way ANOVA and ANCOVA analysis showing the effects of temperature (temp), volumetric water content (VWC), and family on growth and xylem development.

**Table S3.** Summary of Three-Way of ANOVA analyses per date showing the effects of temperature (temp), volumetric water content (VWC), and family on photosynthesis, chlorophyll fluorescence, spectral reflectance, water potential and photosynthetic pigments.

**Table S4.** Post-hoc Tukey pairwise comparisons of the xylem anatomical traits that significantly differed in Fig 6.

