## Supplemental Materials for "Stomatal and xylem plasticity, not growth rate, determines white spruce resilience to warmer and drier climates"

### Methods S1 Installed Sensors in KSR Plots.

Air temperature and humidity was recorded using air sensors above the height of the canopy, approximately 60 cm (Models VP3 and VP4, METER Group, Pullman, WA, USA). Daily mean, minimum and maximum values were calculated for air temperatures. VWC was recorded using soil sensors at a depth of approximately 20 cm and daily mean values were calculated (Models 5TM and TE11, METER Group, Pullman, WA, USA). A correction factor was calculated per plot to correct for sensor drift and aging of the 5TM and TE11 probes that were permanently installed in the experimental plots using manual soil VWC measurements taken with a HydroSense system with the CS620 sensor (Campbell Scientific Inc., Edmonton, AB, Canada; Fig. S5 and S6).

### Method S2 Regression-based estimation of leaf temperature

Leaf temperature could not be measured in 2021 during the experiment due to sensor failure. To address this, daily mean leaf temperatures were reconstructed using a simplified regression approach trained on a corresponding 2020 dataset collected under the same experimental design (Murphy *et al.* 2025). Using the 2020 data, separate ordinary least squares models were fit to predict control leaf temperature from control air temperature, and warming leaf temperature from warming air temperature and heater output. Model performance was evaluated using a single 50/50 train–test split (Figure S7). After validation, the models were refit using the full 2020 dataset and applied to the 2021 environmental data to estimate daily mean leaf temperatures for each treatment (Figure S8). Uncertainty in the predictions was quantified using bootstrapping (2000 re-samples) to generate 95% confidence intervals for control and warming leaf temperatures calculated from 5-day running averages.

### Methods S3 Xylem sample preparation.

Sections were stripped of bark and re-cut into sections of approximately 0.5 cm to optimize paraffin infiltration. Samples underwent fixation with 4% paraformaldehyde overnight at 4˚C, a rinse with 1x PBS on ice, dehydration with ethanol solutions of increasing concentrations (70, 85, 95, 100%), paraffin infiltration with solutions of increasing xylene: ethanol concentrations (33, 50, 66, 100%) and then solutions of increasing paraffin: xylene concentrations (33, 50, 66, 100%) under 60˚C incubation. The samples were then placed transversely into silicon cube molds filled with molten paraffin and hardened. The hardened paraffin blocks were then cut into 14µm thick cross-sections using a Leica SM2000R microtome (Leica Biosystems, Nussloch, Germany). Cross-sections were rinsed with pure xylene, then ethanol solutions of decreasing concentrations (100, 95, 70, 50, 30, 0%) before being stained in a solution (2:1 v/v) of 1% Alcian blue in water and 1% safranin in 50% ethanol for 20 min (Hamann, Smets & Lens 2011). Cross-sections were then rinsed with solutions of increasing ethanol concentrations (0, 30, 50, 70, 95, and 100%), pure xylene, and then mounted onto slides with xylene-based mounting medium

### Methods S4 Primary xylem and derived anatomical traits assessed by ROXAS.

Primary xylem anatomical traits: ring area (RA); total cell number (CNo); cell density (CD; number of cells per $mm^2$); cell lumen area (LA); radial and tangential lumen diameter ($D_{rad}$ and $D_{tan}$); and thickness of radial, tangential, and all cell walls ($CWT_{rad}$, $CWT_{tan}$, and $CWT_{all}$). Derived xylem anatomical traits assessed were: relative wood density (RWD; cell wall area per total cell area), a measure of wood density by cell; hydraulic diameter of an individual cell ($D_h$), correcting for effect of elliptical shape on flow (used for earlywood and latewood means in Fig. **4**; Lewis and Boose, 1995); hydraulically weighted mean diameter at the ring scale ($D_{h,ring}$), a weighted average diameter based on contribution to hydraulic conductance (used for ring means in Fig **S7**; Kolb and Sperry, 1999); theoretical hydraulic conductivity ($K_{h(t)}$), as estimated using Hagen-Poiseuille's law adjusted for elliptical tubes (Nonweiler, 1975); theoretical specific hydraulic conductivity ($K_{s(t);}$ i.e., $K_{h(t)}$ normalized to total xylem area); and cell wall reinforcement ($[t/b]^2$), an index for implosion resistance (Hacke et al., 2001).

**Hacke UG, Sperry JS, Pockman WT, Davis SD, McCulloh KA. 2001.** Trends in wood density and structure are linked to prevention of xylem implosion by negative pressure. *Oecologia.* 126(4): 457–461. https://doi.org/10.1007/s004420100628.

**Hamann T, Smets E, Lens F. 2011.** A comparison of paraffin and resin-based techniques used in bark anatomy. *Taxon*. 60(3): 841–851. https://doi.org/10.1002/tax.603016.

**Kolb KJ, Sperry JS. 1999.** Differences in drought adaptation between subspecies of sagebrush (*Artemisia tridentata*). *Ecol*. 80(7): 2373–2384. https://doi.org/10.2307/176917.

**Lewis AM & Boose ER. 1995.** Estimating volume flow rates through xylem conduits. *Am J Bot* 82(9): 1112–1116. https://doi.org/10.1002/j.1537-2197.1995.tb11581.x

**Nonweiler T. 1975.** Flow of biological fluids through non-ideal capillaries. *Encycl Plant Physiol.* 1: 474–477.

(A)

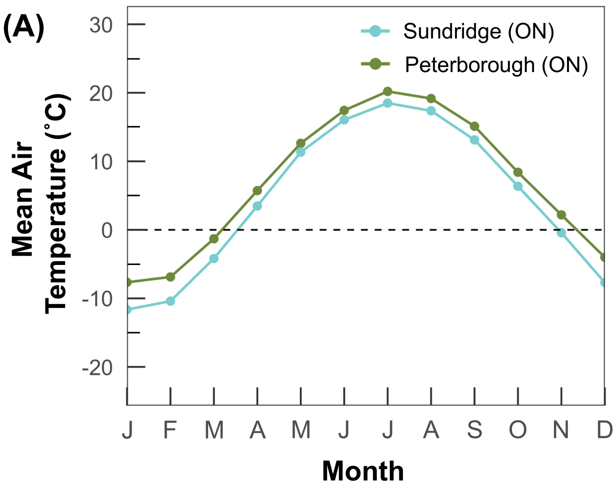


(B)

| | Annual Environmental Data (2000-2020) | | | | | |
|---|---|---|---|---|---|---|
| **Parental Origin Site** | **Min Temp (˚C)** | **Mean Temp (˚C)** | **Max Temp (˚C)** | **Min Rainfall (mm)** | **Max Rainfall (mm)** | **Total Rainfall (mm)** |
| Peterborough, ON | 0.99±0.83 | 6.75±0.79 | 12.52±0.82 | 654.4 | 1118.3 | 862.2±114.0 |
| Sundridge, ON | -1.63±0.83 | 4.33±0.81 | 10.27±0.86 | 784.7 | 1519.4 | 1119.4±163.8 |

**Fig. S1 Seasonal temperatures and annual precipitation for the period 2000-2020 at the parental origin sites for the two families of white spruce.** The fast-growing family was obtained from a cross between two genotypes from Peterborough, Ontario (44°18'17"N, 78°19'12"W). The slow-growing family was obtained from a controlled cross between Sundridge, Ontario (45°46'08"N, 79°23'47"W). (**a**) Average monthly minimum air temperatures; (**b**) table displaying average annual minimum, mean, and max temperatures (± SD), and minimum and maximum annual precipitation. The environmental data was forecasted based on regional air temperature and precipitation interpolated from nearby weather stations and adjusted for elevation and location differentials with regional gradients using the software BioSIM (https://cfs.nrcan.gc.ca/projects/133).

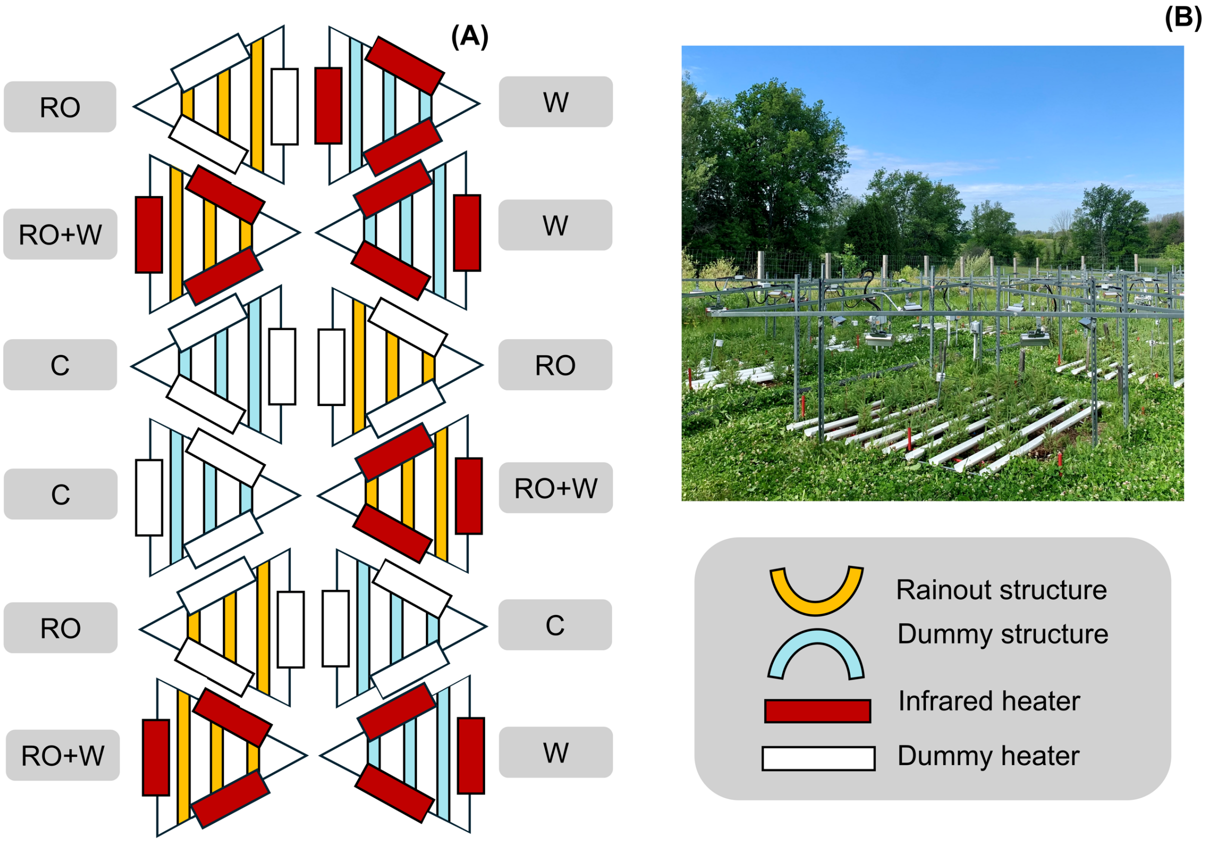


**Fig. S2 The temperature free-air-enhancement and rainout experimental set up at the Koffler Scientific Reserve in Ontario, Canada. (a)** Schematic of the randomly assigned climatic treatments to the twelve plots at the experimental site; **(b)** Plots equipped with infrared heaters and rainout structures. Control, C; Rainout, RO; Warming, W; Rainout combined with warming, RO+W.

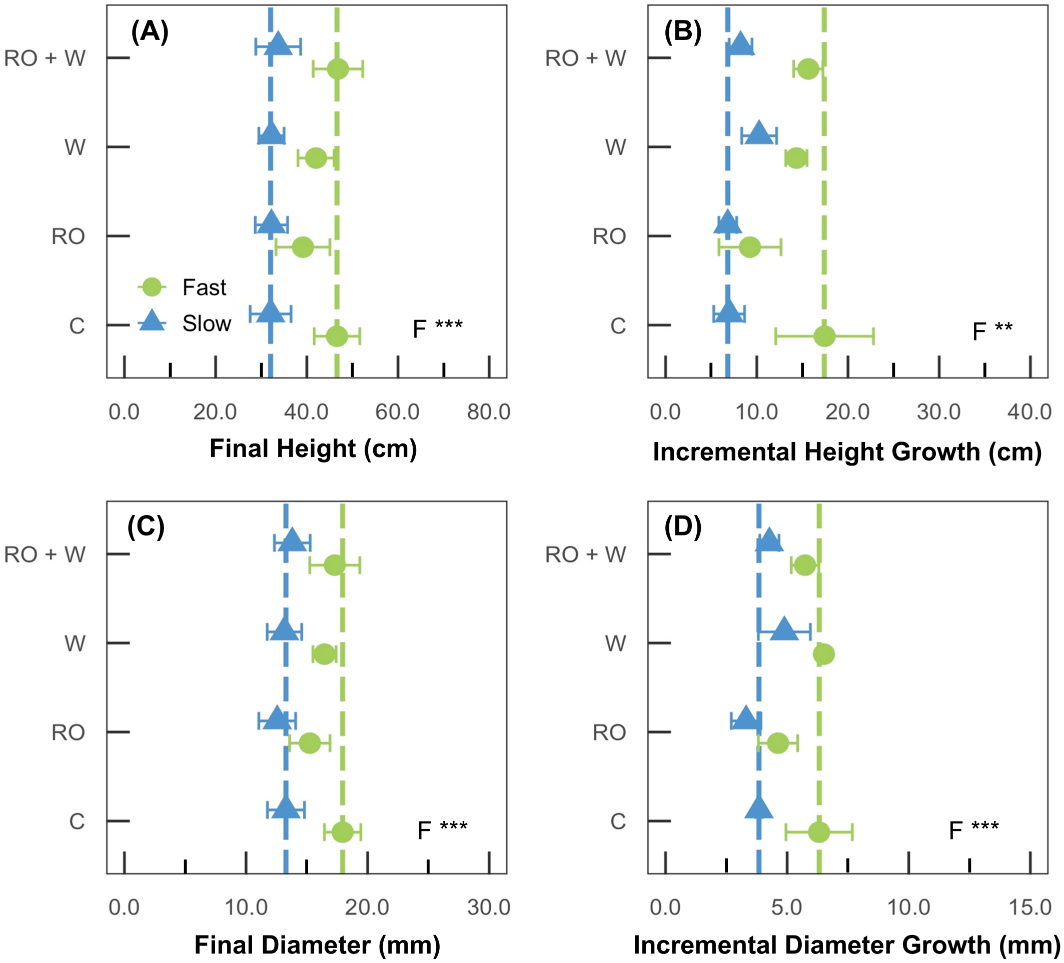


**Fig. S3 The impacts of summer rainout and warming on aboveground growth in a fast- and slow-growing family of white spruce. (a)** total height; **(b)** total diameter; **(c)** difference in seedling height from the start to the end of the experiment (incremental height growth); **(d)** difference in seedling diameter from the start to the end of the experiment (incremental diameter growth). Initial measurement date was May 7 and final measurement date was September 28. Control, C; Rainout, RO; Warming, W; Rainout combined with warming, RO+W. Data points represent the average of three plots, ± SE. Vertical lines represent values for the control seedlings for the fast-growing family (green) and slow-growing family (blue). Temperature (T), volumetric water content (VWC), and family (F) were tested as main effects. Only significant main effects or interactions are noted ($P<0.05$). *$P \leqslant 0.05$, **$P < 0.01$ and ***$P < 0.001$.

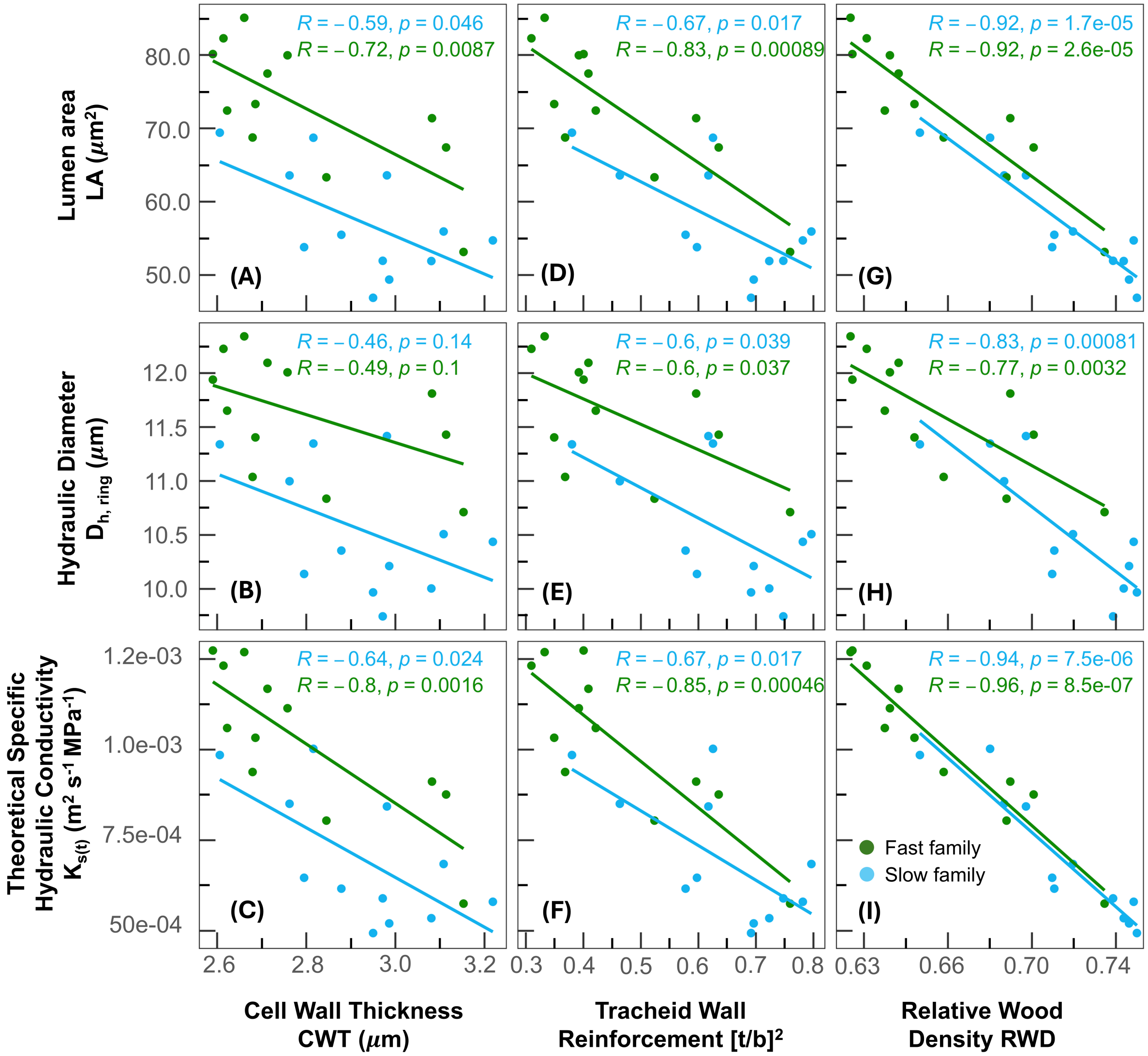


**Fig. S4. Correlations between primary and derived xylem anatomical traits related to hydraulic efficiency and hydraulic safety in a fast- and slow-growing family of white spruce.** Traits shown include: **(a,d,g)** lumen area (LA), **(b,e,h)** hydraulically weighted mean diameter at the ring scale ($D_{h,ring}$), and **(c,f,i)** theoretical specific hydraulic conductivity ($K_{s(t)}$), each plotted against **(a,b,c)** cell wall thickness (CWT), **(d,e,f)** tracheid wall reinforcement ($[t/b]^2$), and **(g,h,i)** relative wood density (RWD; cell wall area per total cell area). Points represent biological averages from across the entire experimental growth ring. Lines show family-specific regressions for fast- (green) and slow-growing (blue) families, with Pearson correlation coefficients (R) and associated p-values shown for each family.

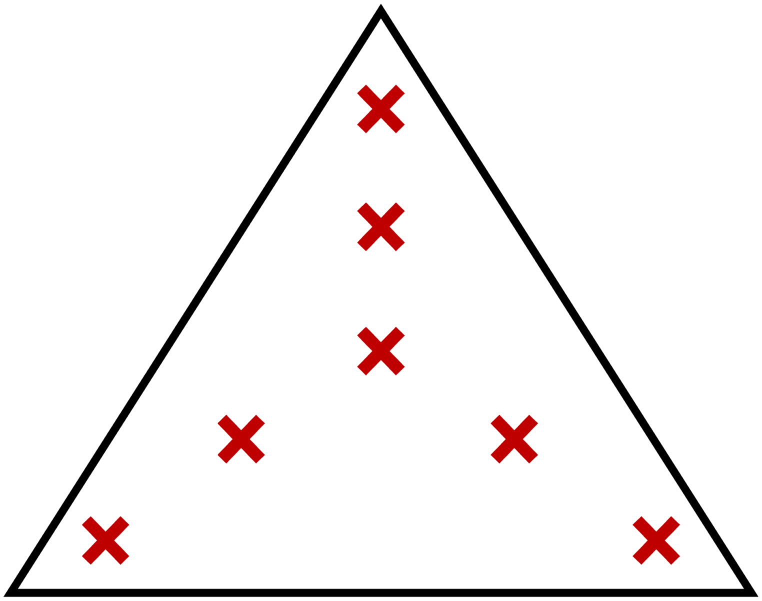

**Fig. S5 Approximate points where manual soil volumetric water content (VWC) was measured and then averaged from in each plot.**

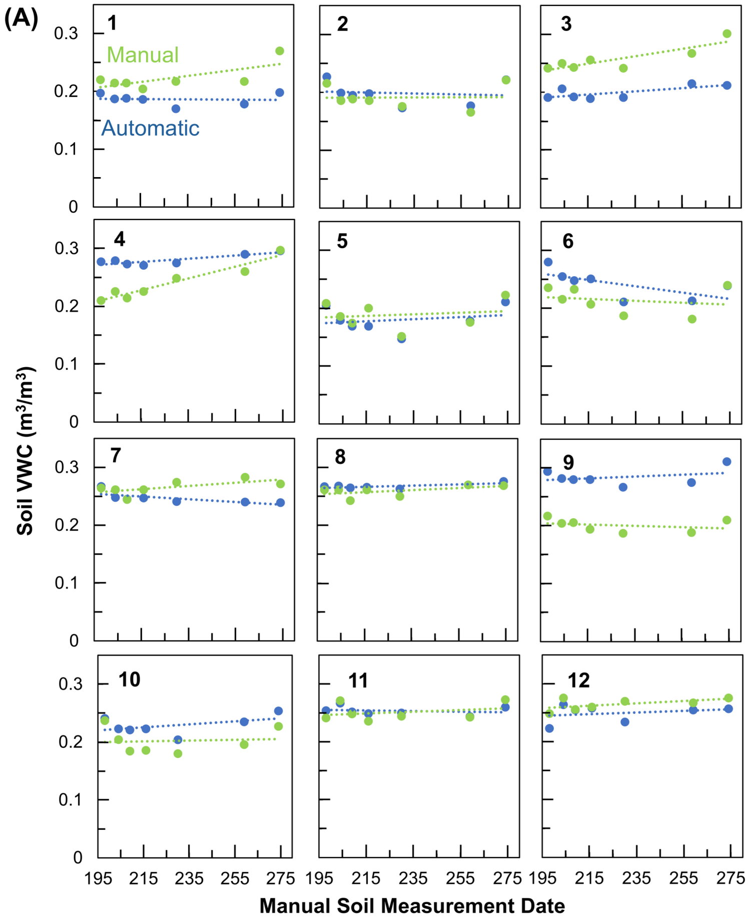


KSR 2021

(B)

| Plot | Corr. Factor |
|---|---|
| 1 | 1.194 |
| 2 | 0.962 |
| 3 | 1.289 |
| 4 | 0.857 |
| 5 | 1.048 |
| 6 | 0.884 |
| 7 | 1.077 |
| 8 | 0.968 |
| 9 | 0.707 |
| 10 | 0.883 |
| 11 | 0.989 |
| 12 | 1.061 |

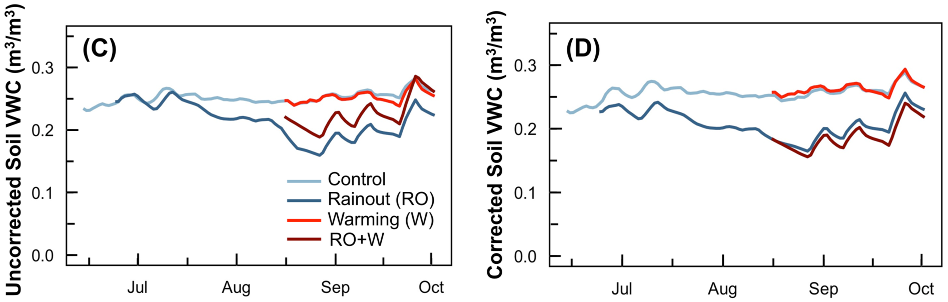


**Fig. S6. Automatic soil sensor corrections applied for the 2021 experimental time frame from June 24 to October 2 at the Koffler Scientific Reserve in Ontario, Canada. (a)** Comparison of the mean manual soil volumetric water content (VWC; green) against the automatic soil sensor (blue) at the same time and date across the experiment within each plot. Plot numbers can be found in the upper left corners of the subplots. **(b)** The calculated correction factors were determined by the ratio of the mean manual soil VWC to the automatic soil VWC points across all time points per plot. **(c)** the 5-day running average of daily soil VWC across each treatment (blue, control; dark blue, rainout; light red, warming; dark red, warming combined with rainout) across the experimental time frame in 2021 before the correction factors were applied. **(d)** the 5-day running average of the corrected daily automatic soil VWC across each treatment across the experimental time frame in 2021 using the correction factors found in Part **(b)** of the figure.

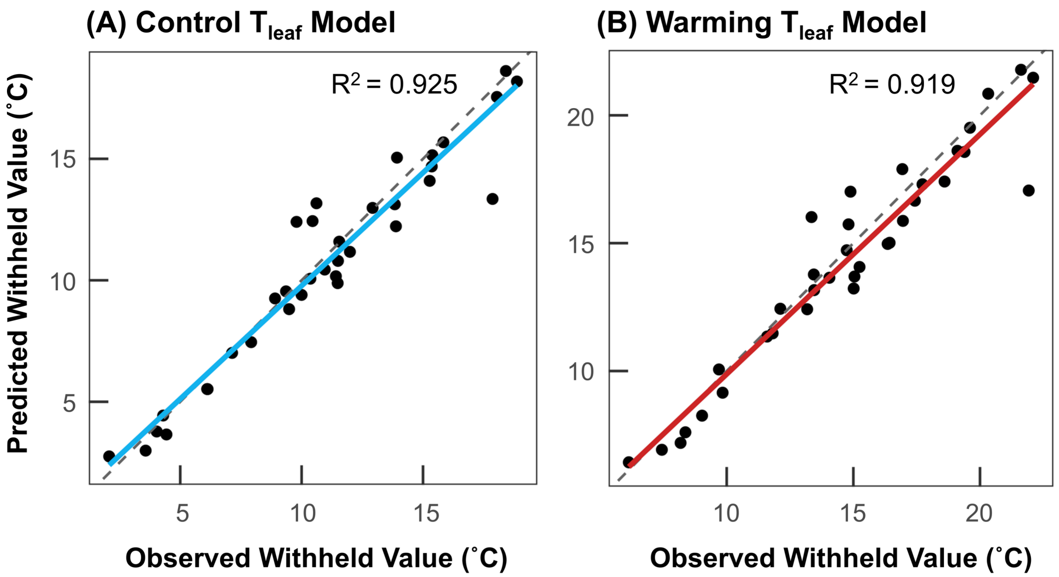


**Fig. S7. Observed versus predicted withheld values from a single 50/50 train–test validation of regression models developed using 2020 data from Murphy *et al.* (2025).** (**A**) Control leaf temperature ($T_{leaf}$) model and (**B**) warming leaf temperature ($T_{leaf}$) model. Points represent withheld observations, solid lines show fitted regression relationships, and dashed lines indicate the 1:1 line. $R^2$ values are shown for each model.

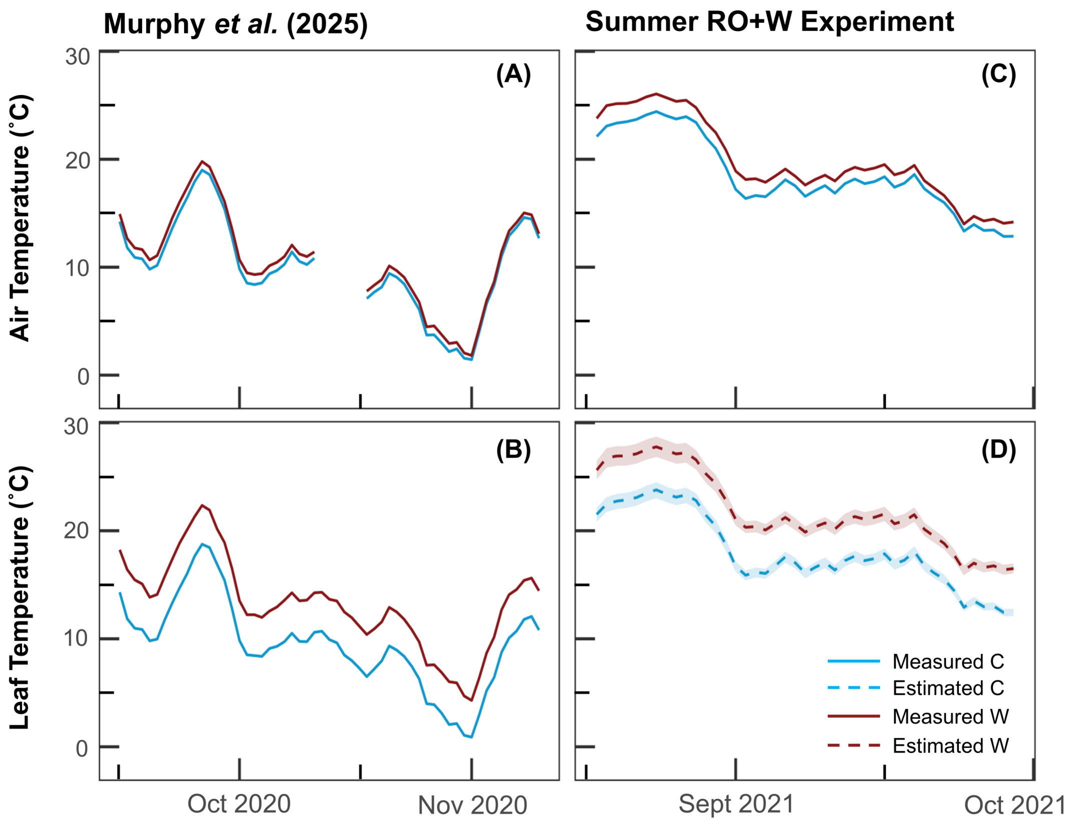


**Fig. S8**. **Comparison of measured and estimated air and leaf temperatures across experiments**. 5-day running averages of air temperature (A, C) and leaf temperature (B, D) for control (blue) and warming (red) treatments. Panels (A) and (B) show measured air and leaf temperatures from the 2020 experiment reported in Murphy *et al.* (2025). Panels (C) and (D) show measured air temperatures and estimated leaf temperatures for the 2021 summer RO+W experiment. Solid lines indicate measured temperatures, and dashed lines indicate estimated leaf temperatures; shaded regions represent 95% confidence intervals around leaf temperature estimates.

**Table S1. Cuvette temperature and relative humidity measured during each measurement campaign using the LI-6400 XT gas exchange system.**

| Date | Treatment | Temperature (˚C) | Relative Humidity (%) | VPD (kPa) | Sample size (n) |
|---|---|---|---|---|---|
| Jun-23 | C | 22.8±0.6 | 56.9±0.6 | 1.55±0.06 | 22 |
| Jul-23 | C | 24.2±0.7 | 55.6±0.3 | 1.70±0.08 | 22 |
| Aug-15 | C | 24.9±0.4 | 54.7±0.5 | 1.78±0.05 | 22 |
| Sept-28 | C | 19.2±0.3 | 56.8±0.4 | 1.22±0.03 | 11 |
| | W | 24.6±0.2 | 47.1±0.6 | 1.81±0.02 | 11 |

*Values represent means±SE.*

**Table S2. Summary of Three-Way ANOVA and ANCOVA analysis showing the effects of temperature (temp), volumetric water content (VWC), and family on growth and xylem development.**

| | | Temp | | VWC | | Family | | Temp x VWC | | Temp x Family | | VWC x Family | |
|---|---|---|---|---|---|---|---|---|---|---|---|---|---|
| | **Variable** | **F** | **P** | **F** | **P** | **F** | **P** | **F** | **P** | **F** | **P** | **F** | **P** |
| Growth | Height Over Time | 0.0 | 0.921 | 0.6 | 0.442 | 58.3 | **<0.001** | NA | NA | NA | NA | NA | NA |
| | Diameter Over Time | 1.3 | 0.263 | 1.3 | 0.252 | 34.9 | **<0.001** | NA | NA | 3.0 | 0.085 | NA | NA |
| | Final Height | 0.1 | 0.751 | 0.0 | 0.947 | 52.7 | **<0.001** | NA | NA | NA | NA | NA | NA |
| | Final Diameter | 0.1 | 0.730 | 0.2 | 0.692 | 37.1 | **<0.001** | NA | NA | NA | NA | NA | NA |
| | Cumulative Height | 1.4 | 0.241 | 1.8 | 0.186 | 13.5 | **0.001** | NA | NA | NA | NA | NA | NA |
| | Cumulative Diameter | 3.4 | 0.090 | 4.0 | 0.067 | 15.0 | **0.002** | NA | NA | NA | NA | NA | NA |
| | Aboveground Biomass | 1.2 | 0.291 | 0.0 | 0.977 | 4.4 | **0.049** | NA | NA | NA | NA | NA | NA |
| | Belowground Biomass | 2.1 | 0.164 | 0.1 | 0.758 | 0.7 | 0.424 | NA | NA | NA | NA | NA | NA |
| | Total Biomass | 1.6 | 0.212 | 0.0 | 0.890 | 3.0 | 0.099 | NA | NA | NA | NA | NA | NA |
| | Root: Shoot | 0.8 | 0.368 | 0.5 | 0.480 | 0.9 | 0.345 | NA | NA | NA | NA | NA | NA |
| Xylem Earlywood | Lumen A | 3.0 | 0.096 | 0.5 | 0.497 | 3.8 | 0.063 | NA | NA | NA | NA | NA | NA |
| | CWT | 2.6 | 0.119 | 0.2 | 0.674 | 0.8 | 0.367 | NA | NA | NA | NA | NA | NA |
| | $D_h$ | 1.9 | 0.184 | 0.4 | 0.552 | 2.5 | 0.123 | NA | NA | NA | NA | NA | NA |
| | $K_{h(t)}$ | 4.2 | 0.052 | 0.2 | 0.684 | 4.2 | 0.005 | NA | NA | NA | NA | NA | NA |
| | RWD | 0.7 | 0.401 | 0.6 | 0.451 | 2.7 | 0.111 | NA | NA | NA | NA | NA | NA |
| | $[t/b]^2$ | 0.5 | 0.480 | 0.7 | 0.409 | 0.9 | 0.345 | NA | NA | NA | NA | NA | NA |
| Xylem Latewood | Lumen A | 24.1 | **<0.001** | 1.4 | 0.253 | 46.1 | **<0.001** | NA | NA | NA | NA | 5.5 | **0.036** |
| | CWT | 2.9 | 0.116 | 0.1 | 0.729 | 4.1 | 0.066 | NA | NA | NA | NA | NA | NA |
| | $D_h$ | 21.9 | **<0.001** | 0.8 | 0.382 | 39.1 | **<0.001** | NA | NA | NA | NA | 5.5 | **0.036** |
| | $K_{h(t)}$ | 22.7 | **<0.001** | 1.1 | 0.319 | 27.3 | **<0.001** | NA | NA | NA | NA | NA | NA |
| | RWD | 3.0 | 0.109 | 0.1 | 0.729 | 32.1 | **<0.001** | NA | NA | NA | NA | NA | NA |
| | $[t/b]^2$ | 0.0 | 0.982 | 0.1 | 0.728 | 31.3 | **<0.001** | NA | NA | NA | NA | NA | NA |

Note: For the ANCOVAs, variables were estimated as Variable ~ Min_Temp + Photoperiod + Temp * VWC * Family + (1|Plot) + (1|ID). For the ANOVAs, variables were estimated as Variable ~ Temp * VWC * Family + (1|Plot). For each parameter, the best fit model including temperature, VWC, family and their interactions were chosen according to the lowest AIC. NA is used to denote a factor or interaction not included in the best-fit model. Temperature, VWC, and family are categorical fixed factors. Plot was included as a random factor. $K_h$ values were log-transformed prior to analysis to improve numerical stability and interpretability due to their very small magnitude.

**Table S3. Summary of Three-Way of ANOVA analyses per date showing the effects of temperature (temp), volumetric water content (VWC), and family on photosynthesis, chlorophyll fluorescence, spectral reflectance, water potential and photosynthetic pigments.**

| | | | Family | | VWC | | Temp | | Temp x VWC | | Temp x Family | | VWC x Family | |
|---|---|---|---|---|---|---|---|---|---|---|---|---|---|---|
| **Dates** | **Variables** | | **F** | **P** | **F** | **P** | **F** | **P** | **F** | **P** | **F** | **P** | **F** | **P** |
| Jun 23 | Gas Exchange | $A_{net}$ | 0.2 | 0.648 | NA | NA | NA | NA | NA | NA | NA | NA | NA | NA |
| | | $g_s$ | 2.0 | 0.181 | NA | NA | NA | NA | NA | NA | NA | NA | NA | NA |
| | | iWUE | 9.6 | **0.011** | NA | NA | NA | NA | NA | NA | NA | NA | NA | NA |
| | | E | 2.3 | 0.143 | NA | NA | NA | NA | NA | NA | NA | NA | NA | NA |
| | | $R_D$ | 0.0 | 0.987 | NA | NA | NA | NA | NA | NA | NA | NA | NA | NA |
| | Chl Fluo | $F_v/F_m$ | 1.8 | 0.193 | NA | NA | NA | NA | NA | NA | NA | NA | NA | NA |
| | | $\phi_{PSII}$ | 1.0 | 0.331 | NA | NA | NA | NA | NA | NA | NA | NA | NA | NA |
| | | $\phi_{NPQ}$ | 0.0 | 0.889 | NA | NA | NA | NA | NA | NA | NA | NA | NA | NA |
| | | $\phi_{f,D}$ | 0.4 | 0.546 | NA | NA | NA | NA | NA | NA | NA | NA | NA | NA |
| | | 1-qP | 0.1 | 0.722 | NA | NA | NA | NA | NA | NA | NA | NA | NA | NA |
| | Spectral Reflectance | PRI | 2.0 | 0.178 | NA | NA | NA | NA | NA | NA | NA | NA | NA | NA |
| | | CCI | 0.0 | 0.834 | NA | NA | NA | NA | NA | NA | NA | NA | NA | NA |
| | | WI | 50.1 | **<0.001** | NA | NA | NA | NA | NA | NA | NA | NA | NA | NA |
| | | NDVI | 0.6 | 0.461 | NA | NA | NA | NA | NA | NA | NA | NA | NA | NA |
| | Water Potential | | 2.0 | 0.184 | NA | NA | NA | NA | NA | NA | NA | NA | NA | NA |
| Jul 23 | Gas Exchange | $A_{net}$ | 2.0 | 0.168 | 0.8 | 0.382 | NA | NA | NA | NA | NA | NA | NA | NA |
| | | $g_s$ | 1.4 | 0.269 | 1.0 | 0.330 | NA | NA | NA | NA | NA | NA | NA | NA |
| | | iWUE | 1.6 | 0.221 | 1.5 | 0.238 | NA | NA | NA | NA | NA | NA | 5.6 | **0.036** |
| | | E | 0.9 | 0.354 | 0.0 | 0.961 | NA | NA | NA | NA | NA | NA | NA | NA |
| | | $R_D$ | 0.1 | 0.721 | 2.5 | 0.142 | NA | NA | NA | NA | NA | NA | NA | NA |
| | Chl Fluo | $F_v/F_m$ | 2.3 | 0.158 | 1.5 | 0.238 | NA | NA | NA | NA | NA | NA | NA | NA |
| | | $\phi_{PSII}$ | 0.1 | 0.802 | 0.3 | 0.619 | NA | NA | NA | NA | NA | NA | NA | NA |
| | | $\phi_{NPQ}$ | 0.6 | 0.440 | 2.0 | 0.183 | NA | NA | NA | NA | NA | NA | NA | NA |
| | | $\phi_{f,D}$ | 1.5 | 0.240 | 1.4 | 0.261 | NA | NA | NA | NA | NA | NA | NA | NA |

| | | | | | | | | | | | | | | |
|---|---|---|---|---|---|---|---|---|---|---|---|---|---|---|
| | | 1-qP | 0.0 | 0.878 | 0.1 | 0.732 | NA | NA | NA | NA | NA | NA | NA | NA |
| | Spectral Reflectance | PRI | 10.7 | **0.007** | 0.0 | 0.889 | NA | NA | NA | NA | NA | NA | NA | NA |
| | | CCI | 1.4 | 0.261 | 0.3 | 0.612 | NA | NA | NA | NA | NA | NA | NA | NA |
| | | WI | 15.2 | **0.002** | 0.1 | 0.764 | NA | NA | NA | NA | NA | NA | NA | NA |
| | | NDVI | 0.6 | 0.458 | 0.4 | 0.559 | NA | NA | NA | NA | NA | NA | NA | NA |
| | Water Potential | | NA | NA | NA | NA | NA | NA | NA | NA | NA | NA | NA | NA |
| Aug 15 | Gas Exchange | $A_{net}$ | 3.7 | 0.069 | 0.1 | 0.772 | NA | NA | NA | NA | NA | NA | NA | NA |
| | | $g_s$ | 5.1 | **0.044** | 0.0 | 0.882 | NA | NA | NA | NA | NA | NA | NA | NA |
| | | iWUE | 0.9 | 0.368 | 0.1 | 0.759 | NA | NA | NA | NA | NA | NA | NA | NA |
| | | E | 10.9 | **0.007** | 0.0 | 0.952 | NA | NA | NA | NA | NA | NA | NA | NA |
| | | $R_D$ | 3.8 | 0.064 | 0.7 | 0.405 | NA | NA | NA | NA | NA | NA | NA | NA |
| | Chl Fluo | $F_v/F_m$ | 0.5 | 0.496 | 5.8 | **0.024** | NA | NA | NA | NA | NA | NA | NA | NA |
| | | $\phi_{PSII}$ | 1.7 | 0.220 | 5.1 | **0.044** | NA | NA | NA | NA | NA | NA | NA | NA |
| | | $\phi_{NPQ}$ | 0.9 | 0.355 | 2.4 | 0.150 | NA | NA | NA | NA | NA | NA | NA | NA |
| | | $\phi_{f,D}$ | 0.0 | 0.868 | 2.6 | 0.123 | NA | NA | NA | NA | NA | NA | NA | NA |
| | | 1-qP | 0.5 | 0.501 | 2.7 | 0.123 | NA | NA | NA | NA | NA | NA | NA | NA |
| | Spectral Reflectance | PRI | 4.9 | **0.046** | 0.1 | 0.715 | NA | NA | NA | NA | NA | NA | NA | NA |
| | | CCI | 0.0 | 0.914 | 0.0 | 0.952 | NA | NA | NA | NA | NA | NA | NA | NA |
| | | WI | 6.2 | **0.020** | 0.2 | 0.653 | NA | NA | NA | NA | NA | NA | NA | NA |
| | | NDVI | 0.5 | 0.504 | 0.1 | 0.786 | NA | NA | NA | NA | NA | NA | NA | NA |
| | Water Potential | | 0.3 | 0.576 | 1.2 | 0.279 | NA | NA | NA | NA | NA | NA | NA | NA |
| | Pigments | Tot $Chl^{-1}$ | 0.8 | 0.396 | 5.7 | **0.034** | NA | NA | NA | NA | NA | NA | 5.7 | **0.034** |
| | | Car $Chl^{-1}$ | 0.0 | 0.988 | 0.1 | 0.740 | NA | NA | NA | NA | NA | NA | NA | NA |
| | | Lut $Chl^{-1}$ | 6.2 | **0.020** | 3.7 | 0.068 | NA | NA | NA | NA | NA | NA | 4.3 | 0.050 |
| | | Zea $Chl^{-1}$ | 12.1 | **0.005** | 1.5 | 0.245 | NA | NA | NA | NA | NA | NA | NA | NA |
| | | VAZ $Chl^{-1}$ | 0.0 | 0.847 | 0.1 | 0.755 | NA | NA | NA | NA | NA | NA | NA | NA |
| | | DEPS | 8.5 | **0.013** | 2.9 | 0.117 | NA | NA | NA | NA | NA | NA | NA | NA |
| Sept 28 | Gas Exchange | $A_{net}$ | 1.3 | 0.278 | 0.9 | 0.373 | 1.5 | 0.252 | NA | NA | NA | NA | NA | NA |
| | | $g_s$ | 4.0 | 0.072 | 0.1 | 0.712 | 18.3 | **0.001** | NA | NA | NA | NA | NA | NA |
| | | iWUE | 1.1 | 0.306 | 0.0 | 0.897 | 29.4 | **<0.001** | 5.9 | **0.024** | NA | NA | NA | NA |
| | | E | 4.7 | 0.056 | 0.2 | 0.680 | 0.9 | 0.376 | NA | NA | NA | NA | NA | NA |
| | | $R_D$ | 0.0 | 0.908 | 0.4 | 0.547 | 13.4 | **0.004** | 11.7 | 0.006 | NA | NA | NA | NA |

| | | | | | | | | | | | | | |
|---|---|---|---|---|---|---|---|---|---|---|---|---|---|
| Chl Fluo | $F_v/F_m$ | 0.3 | 0.601 | 0.8 | 0.380 | 2.1 | 0.164 | NA | NA | NA | NA | NA | NA |
| | $\phi_{PSII}$ | 8.5 | **0.013** | 0.7 | 0.435 | 1.3 | 0.279 | NA | NA | NA | NA | NA | NA |
| | $\phi_{NPQ}$ | 5.8 | **0.033** | 0.2 | 0.643 | 0.0 | 0.837 | NA | NA | NA | NA | NA | NA |
| | $\phi_{f,D}$ | 0.7 | 0.399 | 0.8 | 0.384 | 14.3 | **0.001** | NA | NA | NA | NA | 9.1 | **0.006** |
| | 1-qP | 5.6 | **0.035** | 1.1 | 0.315 | 1.1 | 0.322 | NA | NA | NA | NA | 13.9 | **0.003** |
| Spectral Reflectance | PRI | 0.4 | 0.566 | 0.6 | 0.470 | 0.3 | 0.565 | NA | NA | NA | NA | 15.6 | **0.002** |
| | CCI | 3.7 | 0.076 | 0.1 | 0.730 | 1.1 | 0.311 | NA | NA | NA | NA | 6.6 | **0.024** |
| | NDVI | 0.8 | 0.391 | 0.1 | 0.783 | 0.1 | 0.802 | NA | NA | NA | NA | 9.0 | **0.011** |
| Water Potential | | 0.6 | 0.441 | 0.6 | 0.444 | 5.1 | **0.043** | NA | NA | NA | NA | NA | NA |
| Pigments | Tot $Chl^{-1}$ | 10.2 | **0.004** | 3.5 | 0.073 | 2.7 | 0.112 | NA | NA | NA | NA | NA | NA |
| | Car $Chl^{-1}$ | 8.1 | **0.009** | 2.1 | 0.158 | 1.5 | 0.223 | NA | NA | NA | NA | NA | NA |
| | Lut $Chl^{-1}$ | 1.7 | 0.203 | 1.3 | 0.260 | 0.0 | 0.846 | NA | NA | NA | NA | NA | NA |
| | Zea $Chl^{-1}$ | 3.0 | 0.110 | 0.6 | 0.461 | 5.9 | **0.032** | NA | NA | NA | NA | NA | NA |
| | VAZ $Chl^{-1}$ | 6.4 | **0.019** | 1.9 | 0.180 | 2.7 | 0.115 | NA | NA | NA | NA | NA | NA |
| | DEPS | 1.2 | 0.296 | 1.5 | 0.244 | 4.0 | 0.068 | NA | NA | NA | NA | NA | NA |

Note: Variables were estimated as Variable ~ Temp * VWC * Family + (1|Plot). For each parameter, the best fit model including temperature, VWC, family and their interactions were chosen according to the lowest AIC. NA is used to denote a factor or interaction not included in the best-fit model. Temperature, VWC, and family are categorical fixed factors. Plot was included as a random factor.

**Table S4. Post-hoc Tukey pairwise comparisons of the xylem anatomical traits that significantly differed in Fig 6.**

| Pairwise Comparisons | | Latewood | | | | |
|---|---|---|---|---|---|---|
| | | LA | $D_h$ | $K_{h(t)}$ | $[t/b]^2$ | RWD |
| C Slow | W Slow | | | | | |
| C Slow | RO Slow | | | | | |
| C Slow | RO+W Slow | | | | | |
| C Slow | C Fast | | | | | |
| C Slow | W Fast | | | | | |
| C Slow | RO Fast | | | | | |
| C Slow | RO+W Fast | | | | | |
| W Slow | RO Slow | | | | | |
| W Slow | RO+W Slow | | | | | |
| W Slow | C Fast | | | | | |
| W Slow | W Fast | | | | | |
| W Slow | RO Fast | | | | | |
| W Slow | RO+W Fast | | | | | |
| RO Slow | RO+W Slow | | | | | |
| RO Slow | C Fast | | | | | |
| RO Slow | W Fast | | | | | |
| RO Slow | RO Fast | | | | | |
| RO Slow | RO+W Fast | | | | | |
| RO+W Slow | C Fast | | | | | |
| RO+W Slow | W Fast | | | | | |
| RO+W Slow | RO Fast | | | | | |
| RO+W Slow | RO+W Fast | | | | | |
| C Fast | W Fast | | | | | |
| C Fast | RO Fast | | | | | |
| C Fast | RO+W Fast | | | | | |
| W Fast | RO Fast | | | | | |
| W Fast | RO+W Fast | | | | | |
| RO Fast | RO+W Fast | | | | | |

| >0.05 | 0.05-0.01 | 0.01-0.001 | <0.001 |
|---|---|---|---|

C, control; RO, rainout; W, warming; RO+W, rainout combined with warming; LA, lumen area; $D_h$, hydraulic diameter; $K_{h(t)}$, theoretical hydraulic conductivity; $[t/b]^2$, tracheid wall reinforcement; RWD, relative wood density. $K_{h(t)}$ values were log-transformed prior to analysis as described in Table S2.
